# Adaptation to an acid microenvironment promotes pancreatic cancer organoid growth and drug resistance in a p53-dependent manner

**DOI:** 10.1101/2023.01.02.522472

**Authors:** Arnaud Stigliani, Renata Ialchina, Jiayi Yao, Dominika Czaplinska, Yifan Dai, Henriette Berg Andersen, Robin Andersson, Stine Falsig Pedersen, Albin Sandelin

**Author notes:** Shared first authors (Arnaud Stigliani, Renata Ialchina, Jiayi Yao, Dominika Czaplinska, Yifan Dai, Henriette Berg Andersen). Corresponding and shared last authors: Stine Falsig Pedersen and Albin Sandelin.

## Abstract

The harsh environments in poorly perfused tumor regions have been proposed to select for traits that may drive cancer aggressiveness. Here, we tested the hypothesis that tumor acidosis interacts with driver mutations to exacerbate cancer hallmarks, including drug resistance, in pancreatic cancer. We gradually adapted mouse organoids from normal pancreatic duct (mN) and early PDAC (mP, with KRAS G12V mutation and +/- p53 knockout), from pH 7.4 (physiological level) to 6.7, representing acidic tumor niches. Acid adaptation rewired organoid transcriptional activity, increased viability and, strikingly, increased Gemcitabine- and Erlotinib resistance. Importantly, this response only occurred in organoids expressing wild-type p53 and was most pronounced when acid-adapted cells were returned to physiological pH (mimicking increased perfusion or invasion). While the acid adaptation transcriptional change was overall not highly similar to that induced by drug adaptation of the organoids, acid adaptation induced expression of cytidine deaminase (*Cda*) and ribonucleotide reductase regulatory subunit M2 (*Rrm2*), both associated with Gemcitabine resistance, and inhibition of these proteins partially restored Gemcitabine sensitivity. Thus, adaptation to the acidic tumor microenvironment increases drug resistance even after cells leave this niche, and this is in part dependent on acid-adaptation-induced transcriptional upregulation of *Cda* and *Rrm2*.

## INTRODUCTION

Pancreatic ductal adenocarcinoma (PDAC) is currently the fourth most frequent cause of cancer-related death worldwide, with a 5-year survival rate of 10% (Siegel et al., 2020) and estimated to become the second leading cause of cancer-related death in Europe and the US by 2030 (Rahib et al., 2021). Due to the lack of specific symptoms and screening tools, about 80% of PDAC cases are already metastatic and inoperable at time of diagnosis (Klein et al., 2013; Sanabria Mateos and Conlon, 2016). Furthermore, existing chemotherapeutic treatments, such as the combination of the epidermal growth factor (EGF) receptor inhibitor Erlotinib and the cytotoxic pyrimidine analog Gemcitabine, only marginally prolong patient survival (Kelley and Ko, 2008; Ma et al., 2010; Moore et al., 2007). This in large part reflects extensive treatment resistance, driven by a combination of multiple biological mechanisms, including genetic instability, a dense, hypoxic and acidic stroma, metabolic changes, and immune suppression (Beatty et al., 2021; Yu et al., 2021). Therefore, to improve diagnosis and treatment options, a better understanding of the mechanisms driving PDAC early resurgence after treatment, and the connected development of drug resistance is necessary.

While the mutational landscape of pancreatic cancer is highly complex and heterogeneous, sequential driver mutations, starting with activating *KRAS* mutations (96% of patient tumors), followed by mutations of *Trp53* (41%)*, SMAD4* and *CDKN2A genes* are thought to be key processes in early pancreatic cancer development (Cicenas et al., 2017; Kanda et al., 2012; Sun et al., 2020). However, despite DNA sequencing of large cohorts of PDAC patient samples (Cancer Genome Atlas Research Network, 2017; Waddell et al., 2015), genetic data alone has so far had limited correlation with either molecular phenotypes or disease specifics beyond general PDAC classification. Since many of these samples were from highly progressed cancer stages, a promising avenue is to study PDAC mechanisms in models recapitulating early cancer development that also can link genetic state with other cancer-driving features, such as the tumor microenvironment (TME) (Cicenas et al., 2017).

Illustrating this point, one of the most frequent *Trp53* (also known as p53) mutations in pancreatic cancer is R273H which renders DNA-binding of p53 pH sensitive (Muller and Vousden, 2013; White et al., 2017). This is particularly intriguing because of the importance of TME acidosis for cancer aggressiveness (Blaszczak and Swietach, 2021; Boedtkjer and Pedersen, 2020; Corbet and Feron, 2017). As tumors grow and are increasingly limited by insufficient vascularization, the highly metabolically active cancer cells generate niches with varying levels of extracellular acidosis, usually ranging from around pH 6.5 - pH 7.0 in various tumor types (Boedtkjer and Pedersen, 2020; Vaupel et al., 1989). Such environments, which may or may not overlap with other stressful conditions such as hypoxia and low nutrient levels, are often toxic to normal cells (Cruz-Monserrate et al., 2014; Feig et al., 2012). In contrast, cancer cells which survive these challenging conditions and become adapted to growth in the acidic TME, can become highly aggressive and exhibit traits such as increased invasiveness (Boedtkjer and Pedersen, 2020; Corbet et al., 2020). While environmental acidosis is known to contribute to treatment resistance in solid tumors, this has been ascribed mainly to trapping of weak base chemotherapeutics in the acidic microenvironment (Trédan et al., 2007), and it is unknown whether treatment resistance is an intrinsic feature of acid-adapted tumor cells. This is of particular interest in pancreatic cancer, where the physiology of this organ may already predispose ductal epithelial cells to increased fitness under acidic conditions (for a discussion, see (Pedersen et al., 2017)). At present, there is little data on how acid adaptation changes key phenotypes of cancer cells, including PDAC cells. We recently showed that p53 knockout (KO) exacerbated aggressive growth induced by acid adaptation of Panc02 pancreatic cancer cells in certain extracellular matrix (ECM) conditions (Czaplinska et al., 2022). However, how synergy between acidosis and driver mutations impacts PDAC treatment resistance is fully unknown. Identification of how these two essential traits of cancers interact may lead to novel mechanistic insight and could reveal new targetable vulnerabilities.

Here, using a combination of gradual external pH adaptation protocols, functional assays, RNA sequencing and pharmacological interventions, we show that acid adaptation of PDAC organoids rewires their phenotype toward more aggressive, cancer-associated traits that are highly relevant for treatment resistance, including increased viability and resistance to the DNA synthesis inhibitor Gemcitabine and the epidermal growth factor receptor (EGFR) inhibitor Erlotinib. Crucially, we show that only organoids with wild type Trp53 genotype developed the specific phenotype. We show that the transcriptional state induced by acid adaptation only to a small degree overlaps with that of drug-adapted organoids, suggesting that these two stressors drive cells towards drug resistance through largely independent trajectories. For acid-adapted organoids, a large part of the increased survival following drug treatment stems from the overall increase in viability gained through the adaptation process. However, we also show that a few key drug metabolism genes such as *Cda* and *Rrm2* are upregulated in acid-adapted organoids, and inhibition of these genes decreased acid-adapted organoid drug resistance substantially more than non-adapted organoids, suggesting that acid adaptation also affects specific drug pathways. Our data strongly suggests that acid adaptation within tumors can select for cellular phenotypes driving cancer treatment resistance.

## RESULTS

### Acid adaptation and subsequent return to physiological pH leads to increased growth

PDAC precursors of the Pancreatic intraepithelial neoplasia (PanIN) type are divided into four sub-categories reflecting the histological grade of the lesion. More than 90% of PanIN-IA lesions harbor the *KRAS* allele encoding the constitutively active variant (KRASG12D) (Kanda et al., 2012). Here, we employed KRASG12D-expressing PanIN-IA-derived mouse organoids, denoted mP4, and organoids derived from normal pancreatic ducts, denoted mN10 (Boj et al., 2015) (Fig. 1A). To study the link between extracellular acidosis and PDAC characteristics, we developed a homozygous *Trp53* knockout from the mP4 line using CRISPR/Cas9, denoted ‘mP4 p53KO’ (Fig. 1A,B). In this line, we stably expressed Trp53R270H, corresponding to the pH sensitive human R273H mutation (White et al., 2017). We denoted this line ‘mP4 p53R273H’. Thus, the four organoid cells represent normal pancreatic duct epithelial cells (mN10), and progressive stages of early PDAC development (Fig. 1A). Next, we subjected these organoid lines to culture at gradually increasing acidity, starting at physiological pH 7.4, defined as ‘starting control’ (Ctrl_w_), and decreasing growth medium pH to 6.7 over 8 weeks (Fig. 1C), followed by culture at this pH up to week 11. We sampled cells at weeks 5, 8 and 11, denoted acid-adapted AA_w5,_ AA_w8_ and AA_w11._ To model the situation in which cancer cells from the acidic tumor core are re-exposed to physiological pH, e.g. during invasion, improved vascularization, or tumor shrinkage during treatment, a subset of AA_w8_ cells were returned from pH 6.7 to pH 7.4 at week 9, cultured at this pH for two weeks, and denoted ‘acid-adapted returned organoids’ (AA→pH7.4). Control organoids were grown for 11 weeks in parallel at pH 7.4 as a time control, denoted Ctrl_w11_.

**Figure 1.**
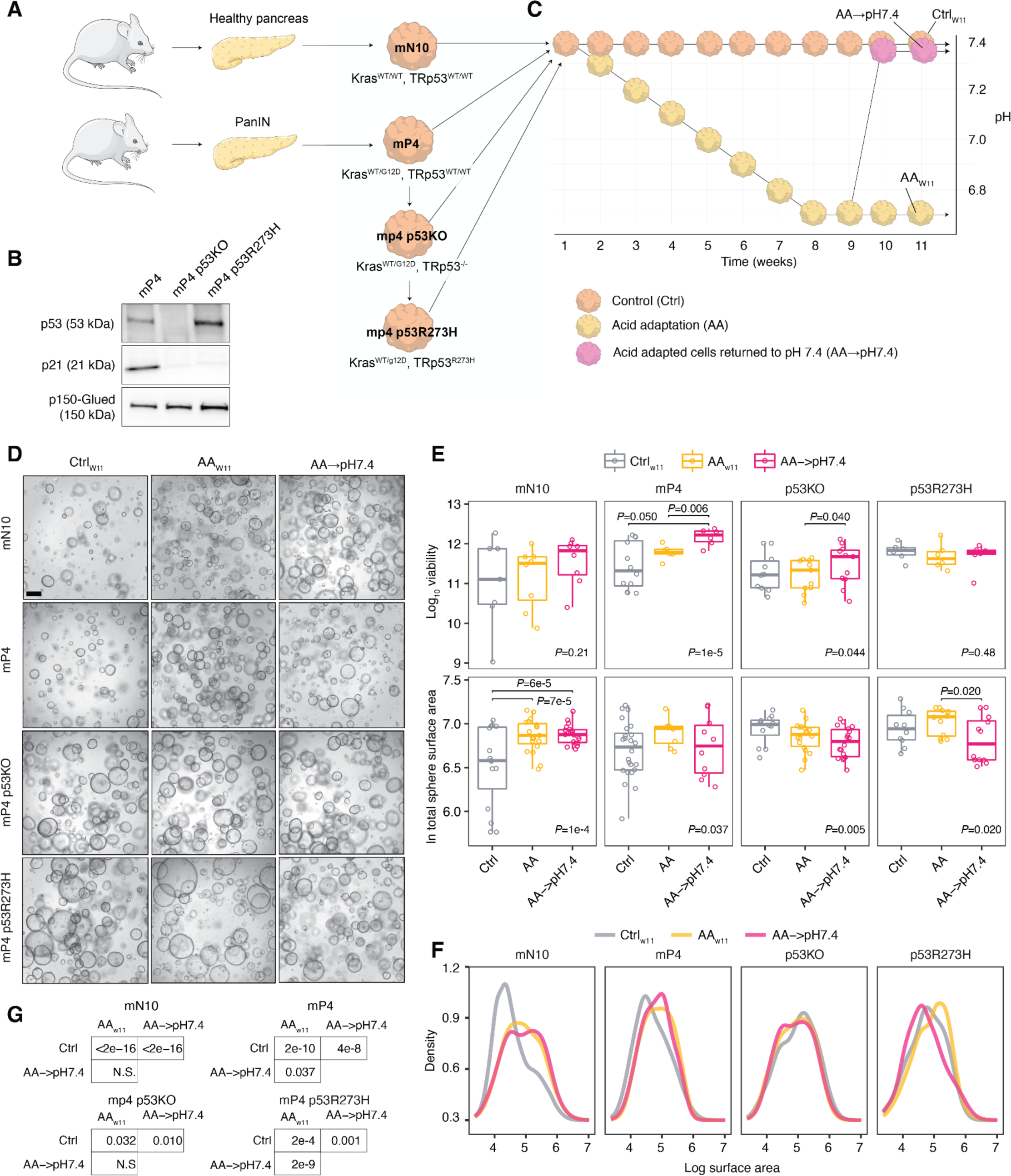
Adaptation to acidic extracellular pH promotes murine pancreatic viability, depending on p53 genotype. **A: Graphical overview of organoid lines used in the project.** mN10 (Kras^WT/WT^, Trp53^WT/WT^) organoids were isolated from healthy murine pancreatic ducts. mP4 (Kras^WT/G12D^, Trp53^WT/WT^) organoids were harvested from murine pancreatic neoplasia (PanIN) and used to develop, with CRISPR/Cas9, p53-deficient line - mP4 p53KO (Kras^WT/G12D^, Trp53^-/-^). mP4 p53KO organoids were subsequently transfected with a vector carrying the DNA sequence coding for p53R273Hto generate the mP4 p53R273H (Kras^WT/G12D^, Trp53^R237H^) line. **B. Western blot validation of p53 KO.** mP4 organoids (Kras^WT/G12D^, Trp53^WT/WT^) were used to generate p53KO (Kras^WT/G12D^, Trp53^-/-^), using CRISPR/Cas9 gene editing. The figure shows a representative Western blot confirming the knockout of p53 and the reintroduction of p53R273H in the mP4 organoid culture. Note the strong reduction in protein level of the p53 target gene p21 in the p53R273H mutant organoids, consistent with earlier reports (Dong et al., 2007). **C. Experimental design for acid adaptation.** All organoid lines/genotypes were gradually adapted to pH 6.7. For each genotype, we ran three panels in parallel, each of them consisting of three independent replicates: i) Control panel (Ctrl.) - organoids cultured at constant pH of 7.4 for 11 weeks, ii) Acid adaptation panel (AA) - organoids gradually adapted to acidosis by decreasing pH of their culture medium by approximately Δ= -0.125 pH unit every week for nine weeks, iii) acid-adapted organoids returned to 7.4 pH (AA→7.4) to mimic return to alkaline pH during metastatic spread. **D. Representative images of murine pancreatic organoids before and after acid adaptation.** Rows show acid adaptation state, as defined in panel C. Columns show genotype. Scale bar: 400 μm. **E: Adaptation to acidic extracellular pH promotes murine pancreatic viability.** Top row shows the distributions of CellTiterGlo viability values for a given genotype and acid adaptation stage (indicated by color). Dots indicate individual observations. *P* values at the bottom indicate results of one-way Anova tests, while pairwise comparison *P* values between conditions are from Tukey’s test. For the latter, only *P* values <0.05 are shown. Bottom row shows the same analysis for total organoid surface area, assessed by Orgaquant image analysis. **F: Size distributions of organoids following acid adaptation.** Each subplot shows the distribution of organoids sizes (in natural log scale) for a given genotype and acid adaptation state (indicated by color). For statistical analysis of distributions, see panel G. Fig. S1A shows the same data but with confidence estimates. **G: Statistical analysis of changes in organoid size distributions as a function of acid adaptation.** Tables show P-values from Kolmogorov-Smirnov test between organoid distributions (from panel E) for each genotype. NS= *P*>0.05.

Representative images of organoid cultures at week 11 for each condition (AA_w11,_ Ctrl_w11,_ AA→pH7.4) are shown in Fig. 1D. As evident from the images, there were substantial differences in organoid numbers and sizes between conditions. Whereas the CellTiterGlo assay measures net cell viability based on ATP content, increased size of individual organoids could indicate increased proliferation/reduced cell death rates, while a greater number of organoids in response to a treatment may suggest a higher stem cell fraction. However, only a few studies have addressed which phenotypic/morphological parameters of organoids best correlate with the impact of, e.g., drug treatments (Beck et al., 2022; Kim et al., 2020). For each condition, we therefore determined (i) total cell viability using the CellTiterGlo viability assay, and (ii) organoid surface areas using Orgaquant analysis (Kassis et al., 2019). The total surface area (Fig. 1E, bottom) is a proxy for the total number of cells, but we also assessed the distribution of surface areas across organoids in a given sample to see if organoid size changed with treatments and/or genotypes (Fig. 1F) Collectively, these measures therefore provide robust readouts of organoid growth patterns in the different conditions and genotypes. mN10 and mP4 organoids showed an increase in viability at AA_w11_ and even more so at AA→pH7.4 compared to Ctrl_w11,_; in the case of mP4, the change was stronger and highly significant (P =1e-5, one-way Anova) and with very low variance at AA_w11_ and AA→pH7.4 (Fig. 1E, top). The total surface area showed similar patterns for both genotypes, although there was no additional increase at AA→pH7.4, and for mP4, the median total surface area had high variance and appeared unchanged vs. Ctrl. (Fig. 1E, bottom). For both of these genotypes, and especially mN10, organoids were significantly larger on average at AA_w11_ and AA→pH7.4 compared to Ctrl_w11_(*P* <2e-16, two-sided Kolmogorov-Smirnov test, Fig. 1G and Fig. S1A) .

Conversely, while mP4 p53KO acid-adapted organoids, particularly AA→pH7.4, showed an increase in viability compared to Ctrl_w11_ almost as large as that of mn10, mP4 p53R273H organoids showed no viability change across acid adaptation states. On the other hand, the latter genotype had a baseline viability level that was higher than that of all other combinations of organoid genotypes and treatments, except AA→pH7.4 mP4 (Fig. 1E, top). mP4 p53KO organoids decreased slightly in total surface area at AA_w11_ and AA→pH7.4, while mP4 p53R273H organoid size was not significantly affected; however, notably, the baseline sizes of organoids from these two genotypes were higher than those of mP4 and mN10 organoids (Fig. 1E, top). Conversely, both mP4 p53KO and mP4 p53R273H genotypes decreased slightly in overall organoid size distribution at AA→pH7.4: interestingly, mP4 p53R273H showed a slight increase at AA_w11_ (Fig1. F,G). Thus, compared to the parental versions, the two p53-compromised organoids showed a different phenotype where viability and total surface areas were less affected and the organoid size was not increased by acid adaptation.

Collectively these results indicate that long-term exposure to the acidic pH of the TME leads to a rewiring that enables increased viability, total organoid area and organoids sizes, in a manner contingent on intact p53 and, in terms of viability, further increased by the KRAS G12D mutation. Thus, the acidic TME selects for traits that endow the cancer cells with a potential for increased growth/survival, in a manner which is strongly affected by key PDAC driver mutations and which is only realized in full when the environment becomes less aggressive.

### Acid adaptation response strategies are diverse and dependent on genetic makeup

We reasoned that if acid adaptation endows cancer cells with a phenotype that renders them more aggressive, it would be important to characterize the gene expression repertoire of cells across the acid adaptation time course and the return to normal pH. Therefore, we sampled Ctrl_w1_, AA_w5,_ AA_w8,_ AA_w11,_ and AA→pH7.4 organoids and subjected all of these to paired end RNA-seq analysis, using biological triplicates for each time point and treatment (Fig. 2A: Fig. S2A and Table S1). To make analyses comparable across time and genotypes, we initially focused on expression changes relative to Ctrl_w1_ for all subsequent time points, for each genotype. Using such a genotype-specific reference removes the effect of genotype-specific starting states. A heatmap visualization of genes that were significantly differentially expressed (*FDR*<0.05 and |log_2_ fold change (FC)| >0.25, by limma voom (Law et al., 2014), Fig. S2B) vs. Ctrl_w1_ at one or more later time points in at least one genotype showed large expression changes occurring in both directions at every time point, where some expression change patterns were shared across genotypes and others were present only in one (Fig. 2B-C). An analysis of overlaps of significantly differentially expressed genes (by limma voom, *FDR*<0.05, |log2FC| > 0.5) between genotypes for each time point, visualized as bar plots (Fig. 2C, also see Fig. S2B-D), enabled two important observations:

**Figure 2:**
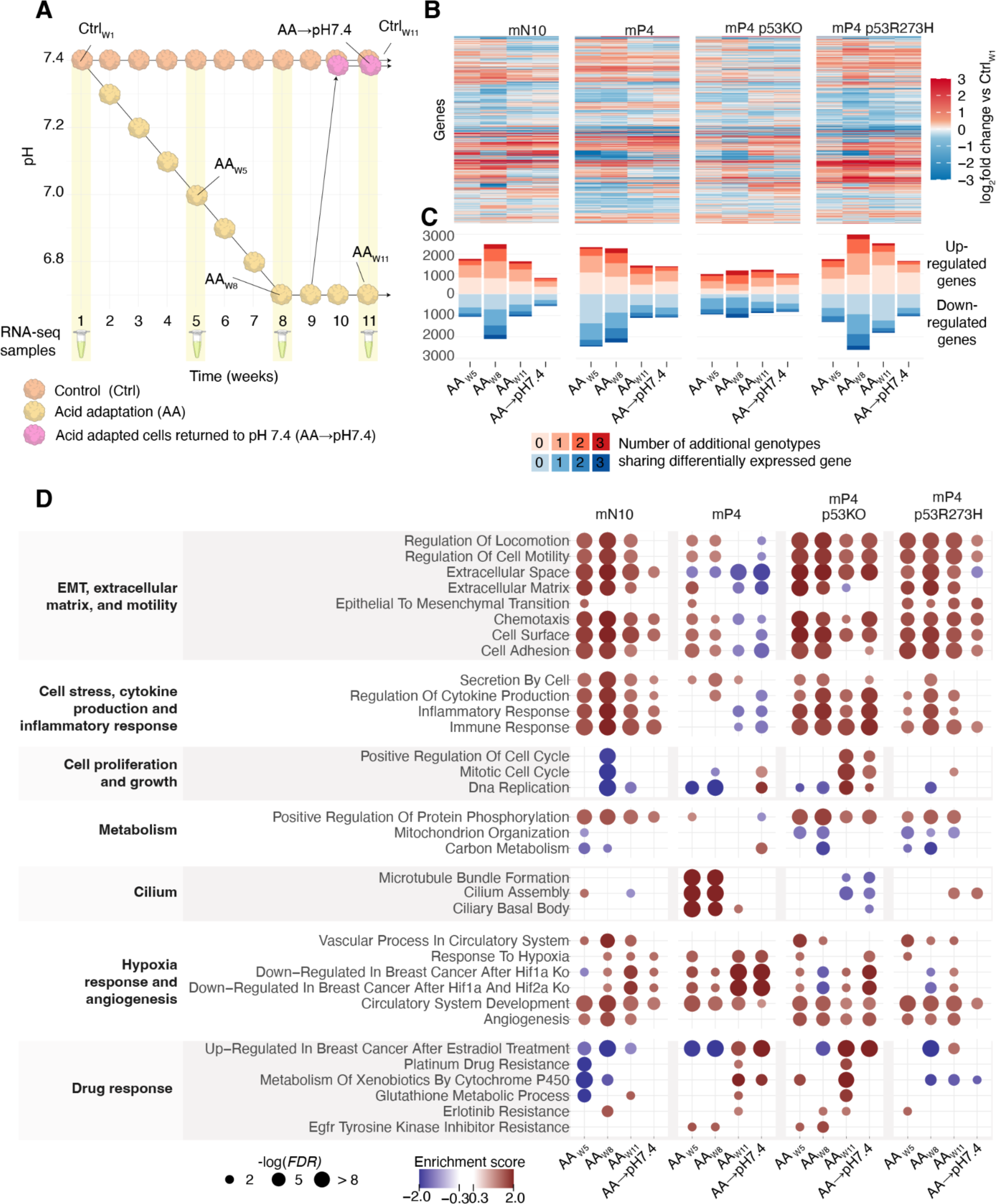
Expression analysis of acid adaption of organoids. **A. Experimental setup.** The plot is organized as in Fig. 1A, with additional indications of which samples were acquired for RNA-seq analysis (marked with yellow highlights and sample names). **B. Expression change vs. Ctrl_w1_ over time.** Expression change is visualized as one heatmap per genotype, where rows were clustered jointly using all genotypes . Rows correspond to genes, columns correspond to adaptation time (AA_w5,8,11_) and subsequent return to physiological pH (AA→pH7.4). Cell colors correspond to average RNA-seq log_2_ FC, comparing respective time points to Ctrl_w1_. **C. Overlap of differentially expressed genes between genotypes across acid adaptation.** Y axis shows the number of differentially expressed genes vs. Ctrl_w1_ for AA_w5,8,11_ and AA→pH7.4), ordered by genotype and time as in the above heatmap and indicated on the X axis. Upregulated genes are shown as positive values in red color tones, down-regulated genes are shown as negative values and in blue color tones. For each genotype and time point, we counted the number of differentially expressed genes that were also differentially expressed in other genotypes at the same time point and in the same direction. These numbers are indicated as color tones in the stacked bar plots, where the deepest color tone indicates genes that are differentially expressed in all genotypes. **D. Gene set enrichment analysis of genes changing expression across the adaptation time course.** For each combination of time point and genotype (shown as columns to the right), we performed a gene set enrichment analysis based on the genes ranked by log_2_FC vs. Ctrl_w1_ as computed in Fig. 2B, and using a large set of gene sets from diverse sources (KEGG, MGsig, GO). Selected gene sets are shown as rows, organized into larger themes as indicated on the left. Dot color shows enrichment or depletion of a given gene (normalized enrichment scores, NES), dot size indicates the associated significance score (-log_10_(*FDR*)).

First, while all genotypes displayed many significantly changing genes at every time point (500-3000 genes per time point): mP4 and mP4 p53R273H organoids had the most, and the mP4 p53KO organoids the least, significantly differentially expressed genes in response to environmental acidosis. mP4 organoids exhibited most changes at early time points during the gradual pH decrease (AA_w5_ and AA_w8_), mP4 p53R273H organoids displayed the largest change once the maximal extracellular acidosis of pH 6.7 was reached (AA_w8_ and AA_w11_) and for mP4 p53KO organoids, the number of changing genes was uniformly distributed across the time points (Fig. 2C).

Second, for a given genotype, roughly half of genes found to be significantly up- or down regulated at any one time point were significantly changing only in that genotype. Conversely, only a small fraction of genes changed significantly in all genotypes (Fig. 2C).

These observations strongly indicate that PDAC cells with different genotypes use at least partially different strategies to adapt and survive in the acidic microenvironment. To obtain insight into such strategies, we performed gene set enrichment analysis of gene sets obtained through GO terms, KEGG (Kanehisa et al., 2021) and Msig databases (Liberzon et al., 2015). Key observations from this analysis are shown in Fig. 2D. In agreement with our previous hypothesis, all genotypes showed overlapping, yet distinct enrichment patterns, where mP4 organoids showed the most distinct pattern of enrichment and depletion. To aid interpretation, functionally related GO terms or gene sets were grouped together. We discuss similarities and differences between genotypes within each such group below.

The mN10, mP4p53KO and mP4p53R273H organoids displayed an upregulation of genes in most of the categories relating to *EMT, ECM and motility* throughout the entire time course, and this persisted in the AA→pH7.4 condition for mP4p53KO and mP4p53R273H organoids. The mP4 organoids stood out by showing weaker upregulation of these terms during the adaptation phase (AA_w5_ and AA_w8_), and a downregulation for many terms at AA_w11_ and for nearly all terms in the AA→pH7.4 condition.

High expression of cytokines and inflammatory response genes is common in PDAC cells even in absence of immune cells (Bellone et al., 2006; Mantovani et al., 2008; Niess et al., 2015). This in large part reflects cancer-associated upregulation of genes such as Interferon-induced protein with tetratricopeptide repeats 3 (IFIT3) (Wang et al., 2020), interleukin-1 (IL-1), chemokine (C-X-C motif) ligand 1 (CXCL1) and other chemokines and chemoattractants, driving a pro-inflammatory phenotype (Somerville et al., 2020). Interestingly, this category of genes was also profoundly altered by acid adaptation. mN10, mP4p53KO and mP4p53R273H organoids displayed comparable enrichment patterns over time for gene sets related to *cell stress, cytokine production and inflammatory response*, with upregulation at all time points, albeit with a generally lower upregulation for mP4p53R273H organoids. This strongly indicates that these genotypes mount an inflammation-like response during the adaptation to acidic pH, and that this state persists in AA→pH7.4 organoids. In contrast, mP4 organoids did not upregulate inflammatory response genes during adaptation, and even down-regulated such genes further at AA_w11_ and AA→ pH7.4(further discussed below).

We observed a general trend for gene sets related to *cell proliferation and growth* to be either unchanged or downregulated in AA_w5_ and AA_w8_ organoids followed by an upregulation in the mP4 and mP4p53KO organoids. In particular, mP4p53KO organoids showed the strongest upregulation at later time points, i.e. in AA_w11,_ and after return to normal pH, i.e. in AA_pH7.4_ organoids. In contrast, mN10 organoids showed substantial downregulation of all terms in these gene sets at 8 weeks. mP4 p53 R273H organoids showed little response in these terms, consistent with their lack of viability change (compare with Fig. 1E). The downregulation of DNA replication terms in all genotypes in AA_w5_ and AA_w8_ is consistent with observation that a substantial number of cells died during these two time points during acid adaptation in all genotypes (see Methods).

All genotypes except mP4 showed an upregulation of *protein phosphorylation*-associated genes, especially at early time points, likely in part reflecting a strong initial signaling response to the stress of being subjected to growth at acidic conditions. In contrast, gene sets related to *mitochondrial organization and carbon metabolism* were initially downregulated in all organoids except mP4, followed by return to levels similar to control values in the AA→pH7.4 condition. This apparently reduced metabolic activity is in line with the generally reduced proliferation and growth in the early time points of acid adaptation, as noted above. Again, the mP4 organoids stood out, with little change in any category, except for a clear increase in carbon metabolism at the AA→pH7.4 condition.

mP4 organoids showed a strong upregulation of cilia-associated genes in the AA_w5_ and AA_w8_ sets while the same sets were downregulated in mP4p53KO organoids at AA_w11_ and AA→pH7.4. This is interesting as ciliogenesis is upregulated in mouse cells resistant to PDAC chemotherapeutics (Jenks et al., 2018) (also see below for analysis of drug resistance of acid-adapted mP4 organoids). Unlike other gene set categories, all genotypes showed roughly similar upregulation of terms relating to *hypoxia response and angiogenesis*, during most or all of the time course. Notably, ‘circulatory systems development’ genes were consistently upregulated in all genotypes throughout the time course, whereas terms strongly related to hypoxia response were most strongly upregulated in mP4 AA_w11_ and AA→pH7.4 organoids. A link between extracellular acidosis and upregulation of genes involved in hypoxia response has previously been reported (Filatova et al., 2016; Mekhail et al., 2004) and this may play a role in the growth increase in these organoids under AA→pH7.4 conditions (compare with Fig. 1D-E).

Strikingly, the mP4 and mP4 p53KO genotypes displayed a unique pattern of upregulation of gene signatures related to drug response in AA_w11_- and also to some extent in AA→pH7.4 - organoids. While mP4 and mP4 p53KO organoids had the highest enrichment (Fig. 2D), we noted that the *Erlotinib resistance* gene profile was enriched in at least one time point for all genotypes. This is interesting in a PDAC context, since Erlotinib, an epidermal growth factor receptor (EGFR) inhibitor, in combination with the DNA synthesis inhibitor Gemcitabine, is a common treatment for metastatic pancreatic cancer (Ma et al., 2010)(Moore et al., 2007).

### Acid-adapted cells develop drug resistance

Based on the GO term analysis above, we hypothesized that acid-adaptation leads to increased drug resistance. To validate this, we treated AA_w11,_ AA→pH7.4 and Ctrl_w11_ organoids from all genotypes for 5 days with 10 or 25 nM of Erlotinib (E), Gemcitabine (G) or both (E+G) and determined viability, number of organoids and total organoid area as described before. We compared this to corresponding data from untreated AA_w11,_ AA→pH7.4 and Ctrl_w11_ organoids (from Fig. 1D).

Fig. 3A shows representative images of organoids following treatments with 10 nM of each drug, alone or in combination. Fig. 3B shows distributions of viability, total organoid surface area and distribution of surface area following 10 nM treatments. Figs. S3A,B show the corresponding images and measurements for 25 nM treatments. For all genotypes except mP4 R273H, regardless of acid adaptation, drug-treated organoids had substantially reduced viability compared to untreated organoids (*P* <=0.006 in all cases except mP4 R273H, one-way Anova) although the effect of drug treatment was attenuated in mP4 p53KO organoids compared to that in mN10 and mP4 organoids. However, for mN10 and mP4, drug treatment of non-acid-adapted untreated organoids induced a more prominent viability reduction compared to drug-treated acid-adapted organoids (Fig. 3B). The same patterns were observed when assessing total organoid surface area, with higher statistical significance(*P* <=1e-6 in all cases except mP4 R273H, one-way Anova, Fig. 3B).

**Figure 3.**
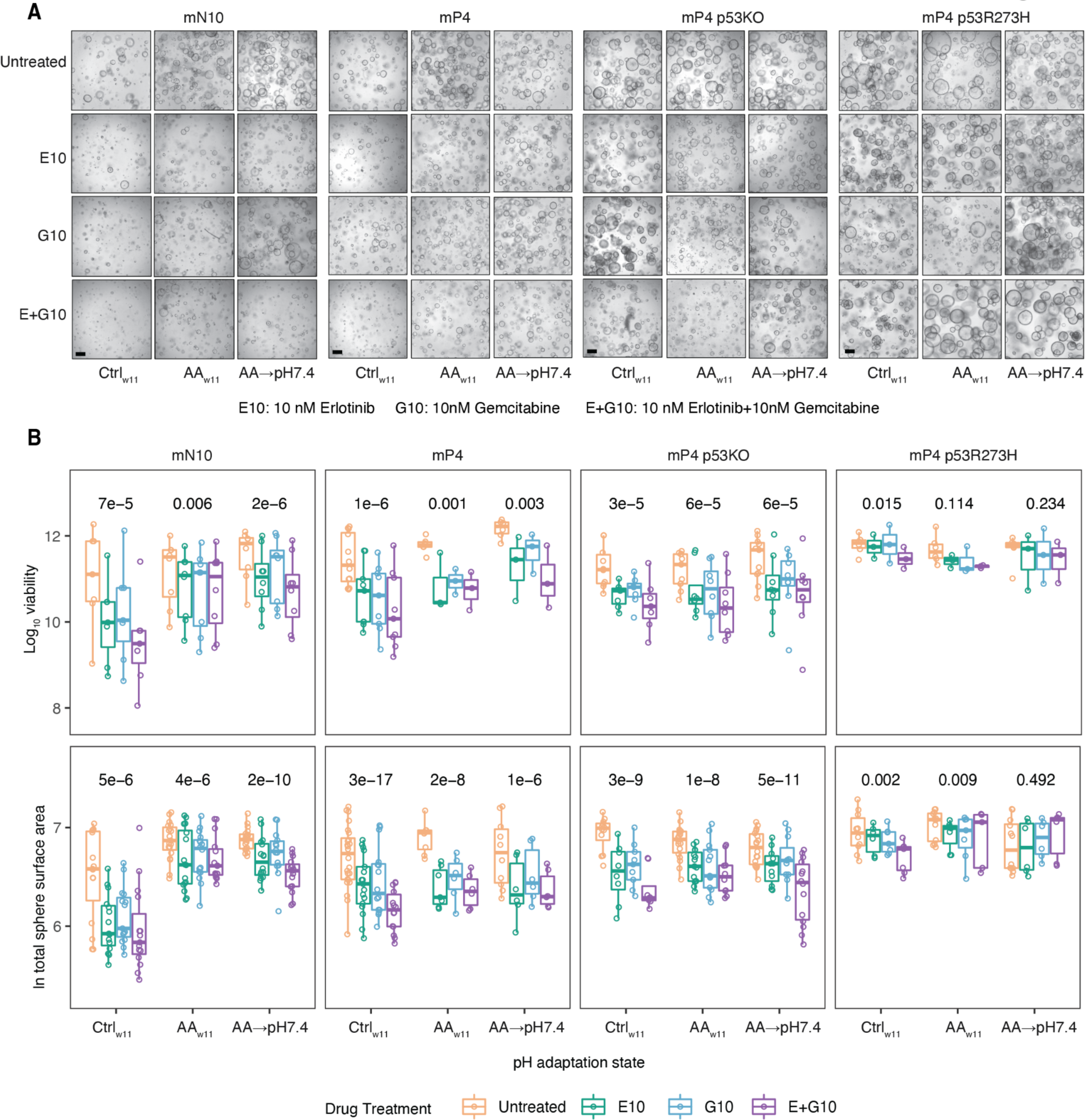
Acid adaptation increases PDAC/PanIN organoid survival after chemotherapy treatment. **A: Representative images of drug-treated and/or acid-adapted organoids.** Columns show acid adaptation conditions, sorted by genotype as in Fig. 1D. Rows show drug treatment. Drug abbreviations are shown beneath images. Only 10 nM treatments are shown (Fig. S3A shows images from 25 nM treatments). Scale bar: 400 μm. **B: Distributions of viability and total organoid surface area following drug treatments.** Boxplots show the distributions of viability and total organoid surface area as in Fig. 1E, but also show data following drug treatments (indicated by color) in respective acid adaptation condition and genotype. For clarity, yellow boxplots (drug-untreated organoids) are based on the same data as in Fig. 1E and included for comparison. One-way Anova *P* values are shown on top, comparing effects of drug treatment in a given acid adaptation and genotype. Fig. S3B shows the above analysis for 25 nM treatments, while Fig. S3C-D summarizes results from linear models for the 10 and 25 nM treatments.

Since we previously established that acid adaptation in these two genotypes by itself led to higher viability and total surface area (Fig. 1D), we hypothesized that the relative increase of viability and total surface area following drug treatments in these genotypes once acid adapted would be the results of two independent effects: i) an increase of viability and total surface area driven by acid adaptation (as shown in Fig. 1D) and ii) a decrease induced by the drug treatment (as shown in Fig. 3B). We tested the assumption of independence by linear models (see Methods), and found that in mP4, mP4 p53KO and mp4 p53R273H organoids, the two effects were largely independent, but in mN10 organoids, there was a significant and positive interaction effect between E+G treatments and AA_w11_ (*P*=0.006, Fig. S3C). This suggests that in mN10 organoids, acid adaptation induced an additional, specific drug resistance effect on top of the baseline increase of viability. This pattern was largely the same, or even stronger, when assessing total organoid surface area: here, the interaction effects were greater and present for almost all drug-acid adaptation combinations (Fig. S3D). Interestingly, these interaction effects were not present for remaining genotypes when assessing viability, while we observed positive interaction effects between most drug-acid adaptation combinations in mP4 p53 organoids when assessing total surface area (Fig. S3C-D).

Treatment using the higher drug concentration, 25 nM, killed the majority of cells in all conditions. Accordingly, the viability and total surface area were significantly decreased (*P* <=0.001 in all cases, one-way Anova, Fig. S3B) for all genotypes including mP4 R273H. Under these conditions, acid adaptation did not seem to confer an increase of viability, and only elicited a weak increase in total surface area for mN10 and mP4 organoids (Fig. S3B-D). While organoid size distribution was largely unaffected by 10 nM treatments, organoids were on average smaller at all 25 nM treatments (Fig. S3D) regardless of genotype and acid adaptation stage.

As a summary, while acid-adapted p53-intact organoids (mP4 and mN10) were sensitive to Erlotinib and Gemcitabine, they showed less decreased viability and total surface area after drug treatment compared to corresponding drug-treated, non-acid-adapted controls. For mP4 cells, this is largely due to the baseline increase in viability that the acid adaptation confers, while in mN10 cells, acid adaptation seems to confer an additional increase in viability when drug-treated. It is interesting to note that mP4 p53KO organoids had a lower overall drug sensitivity baseline, but on the other hand were less affected by acid adaptation state, while mP4 p53R273H organoids were largely unresponsive to both acid adaptation and drug treatments.

### Comparison of acid adaptation and drug adaptation transcriptional change

In the previous analysis, we observed that acid-adapted organoids with intact p53 (mN10 and mP4) survived drug treatments better than corresponding non-acid-adapted organoids. This raised three hypotheses:

First, acid-adapted organoids may survive drug treatments better not through a drug response-specific effect but rather through the general increase in growth (Fig. 1B), as discussed above. An argument for this is that only mN10 organoids had a statistically significant interaction between drug treatment and acid adaptation effects in terms of viability (Fig. 3B).

Second, the acid-adaptation may drive their gene expression to a state similar to that displayed by organoids that acquire drug resistance through actual drug exposure, either over time (drug adaptation) or by acute drug treatment. An argument for this hypothesis is the observed over-representation of drug resistance-associated GO terms in acid-adapted organoids (Fig. 2E).

Chemotherapy resistance in pancreatic cancer may involve changes in drug uptake, metabolism, cell death- or survival signaling, or, often, a combination of these mechanisms (Beatty et al., 2021). Hence, a third hypothesis is that acid-adaptation causes transcriptional rewiring to a state of *bona fide* drug resistance, but not via the same molecular mechanisms as those induced by the Erlotinib and Gemcitabine treatments. Importantly none of these hypotheses are mutually exclusive, and a combination of them could underlie our observations.

To investigate whether acid-adaptation induced a transcriptional state similar to that of adaptation to the drugs *per se*, we exposed mP4 organoids to increasing concentrations of Erlotinib, Gemcitabine or the combination of the two over 12 weeks (Fig. 4A, light blue arrows): we will refer to these as drug-adapted organoids. We chose mP4 organoids for the analysis rather than mN10 since these cells are more PDAC-relevant. Control organoids not subjected to drug treatments were maintained in parallel (Fig. 4A, black arrow). A subset of control organoids were acutely treated with 10 nM Erlotinib, Gemcitabine or both drugs at 12 weeks (Fig. 4A, deep blue arrows): we will refer to these as acutely drug-treated organoids. The drug resistance phenotype was confirmed by treatments with each drug individually or in combination (Fig. S4A-C). To assess the RNA expression changes after drug treatments, we extracted RNA, performed RNA-seq at 12 weeks for each trajectory, and measured expression change as the log_2_ FC ratio between drug treatment and control at week 12 (Fig. 4A).

**Figure 4.**
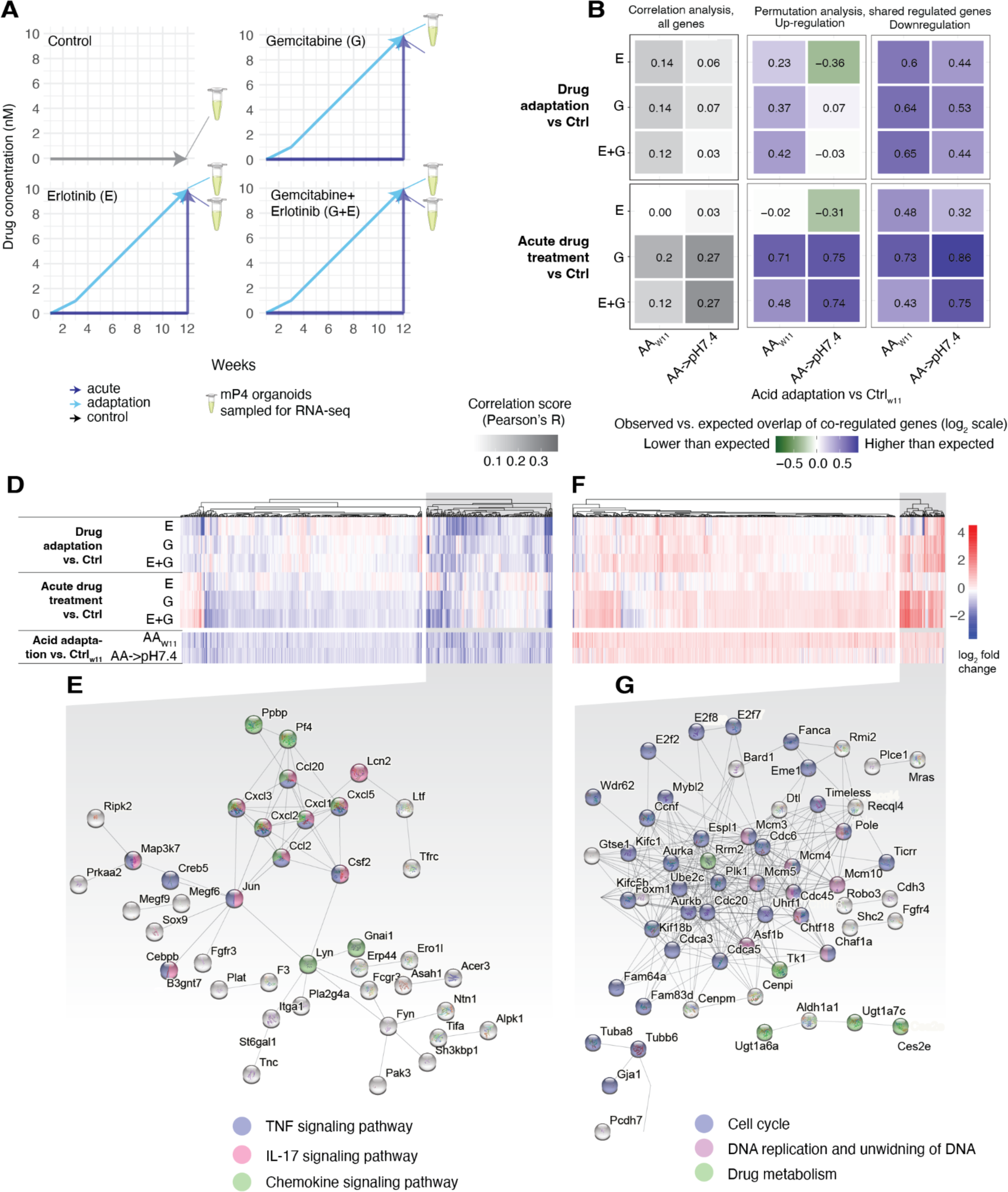
Comparison of acid adaptation and drug treatment gene expression response in mP4 organoids: A: Experimental design of drug treatments of mp4 organoids. Each image shows one experiment where the Y axis shows drug concentration as a function of time in weeks (X axis). The top panel shows the control experiment design, where mp4 organoids are not exposed to drugs but sampled after 12 weeks (black arrow). Remaining panels show treatment with Gemcitabine (G), Erlotinib (E), or the combination (G+E) . In these panels, light blue arrows show adaptation experiments where mP4 organoids were gradually adapted to an increased drug concentration from 0 to 10 nM over 12 weeks. Deep blue arrows show acute treatments, where a subset of control mP4 organoids (black lines) were acutely exposed to 10 nM of respective drug/drug combination at week 12. RNA-seq samples were taken at 12 weeks for each treatment type. **B: Correlation between acid and drug treatment gene expression response.** The heatmap shows pairwise correlation analyses between gene expression responses following drug treatments (rows, split between adaptation and acute treatments as defined in panel A) vs. acid adaptation (columns): gene expression response is defined as the log_2_FC of treatment vs. relevant control RNA-seq expression values as indicated for respective row and column. Cell colors (greyscale) indicate Pearson R values, which are also written out as numbers. **C: Permutation tests of shared up- and down-regulated genes.** For each pairwise analysis, we counted the number of genes which were upregulated in both conditions (log_2_FC >0.25 in both conditions), or downregulated in both conditions (log_2_FC <-0.25 in both conditions). In order to assess whether this number was higher or lower than expected, we made a permutation analysis where gene labels are randomly assigned. The heap map colors show the log_2_(observed vs. expected shared up- or downregulated genes) ratio. Rows and columns as in panel B. **D: heatmap visualization of genes downregulated in acid treatments and >=4 /6 of drug treatments.** Rows indicate treatment vs. controls as defined in panel B. Columns indicate the 627 genes that were upregulated (log_2_FC >0.25) in both acid treatments and at least 4 drug treatments. Cell colors indicate log*_2_*FC gene expression values as defined to the left. Gray background color indicates a subset of genes with particularly strong and downregulation, visualized further in panel E. Columns are clustered only based on drug treatment values. **E: STRING database visualization of strongly connected genes from panel E.** Only the largest cluster is shown. Colors indicate GO term annotations of specific genes. **F: Heatmap visualization of genes upregulated in acid adaptation and >=4/6 of drug treatments.** The image is organized as in panel D, except that the 930 genes that were upregulated (log_2_FC > 0.25) in AA_w11_ and upregulated in at least 4/6 drug treatment experiments were analyzed. **G: STRING database visualization of strongly connected genes from panel F.** The plot is organized as in panel E. Only the largest three clusters found by STRING analysis are shown.

Next, we assessed the pairwise correlation of these drug vs. control log_2_FC to AA_w11_ vs. Ctrl_w11_ and AA→pH7.4 vs. Ctrl_w11_ log_2_FC (left panel of Fig. 4B). These conditions showed at best a modest similarity (mean Pearson’s R value 0.12, max 0.27). We reasoned that while the overall similarity was low, subgroups of genes might show higher similarity, in particular for substantially up-and down-regulated genes in a given pairwise comparison. Therefore, for a given comparison, we counted the number of genes that were upregulated (log_2_FC >0.25) in both treatments, and assessed whether this number was higher or lower from what was expected by chance by permutation analysis. The same method was used to analyze the downregulated genes (log_2_FC < -0.25) in both treatments. Observed vs. expected ratios, expressed at log2 scale, are shown in Fig. 4C. As a summary, while we in most comparisons observed a higher number of jointly regulated genes than expected by chance, this phenomenon was most substantial for downregulated genes and for comparisons involving acute treatments with Gemcitabine or both drugs in combination.

While doing this analysis, we noted that a small subset of genes exhibited similar expression changes across most, if not all, treatments. To explore these, we selected 627 genes that were downregulated (log_2_fc < -0.25) in both AA_w11_ and AA→pH7.4, and in addition downregulated in at least 4/6 drug treatment conditions. A heatmap visualization of these genes (Fig. 4D) showed a subset of genes with particularly strong downregulation (indicated by gray box). A STRING database (Szklarczyk et al., 2019) visualization showed that many of these genes were part of the Tumor necrosis factor (*Tnf*)-, Interleukin-17 (*Il17*)- and chemokine signaling pathways (Fig. 4E), including several chemokine (C-X-C motif) ligand (*Cxcl*) family members *Cxcl2,3 and -5*, Colony Stimulating Factor 2 (*Csf2*), Map3k7 (*Tak1*) and CCAAT/enhancer-binding protein beta (*Cebpb*). This echoes the downregulation of immune response-related genes in AA_w11_ and AA→pH7.4 vs. Ctrl_w1_ for mP4 organoids observed in Fig. 2D. Two other notable downregulated gene clusters were linked to the transcription factor *Jun* and the non-receptor tyrosine kinase *Lyn*, respectively, in the STRING analysis. The *Jun* cluster included receptor-interacting protein kinase (*Ripk*), a mediator of inflammatory cell death signaling (Topal and Gyrd-Hansen, 2021), *Prkaa2*, encoding for AMP-activated protein kinase (*Ampk*), Fibroblast growth factor receptor-3 (*Fgfr3*), and *Sox9*, a key regulator of early pancreas development and a context-dependent oncogene (Panda et al., 2021). The Lyn-related cluster was dominated by regulators of ECM interaction and motility, such as integrin-alpha1 (*Itga1*), the ECM protein Tenascin-C (*Tnc*), and Netrin-1 (*Ntn1*) (Sung et al., 2019).

We used a similar approach for identifying shared upregulated genes, but since the overlap between acid adaptation-up-regulated genes and drug adaptation-up-regulated genes was on average lower than in the analysis above (Fig. 4B-C), we used a more relaxed criterion and selected 930 genes that were upregulated (log2fc > 0.25) in AA_w11_, and upregulated in at least 4 of the 6 drug treatment experiments, and visualized those as a heatmap (Fig. 4F). The heatmap of the expression of these genes showed a cluster of genes with particularly strong upregulation. STRING analysis (Fig. 4G) revealed that many of these genes were highly functionally connected with roles in regulation of cell cycle and DNA unwinding (blue), drug metabolism (green), and DNA replication (pink). Notably, ribonucleotide reductase enzyme subunit M2 (*Rrm2*) was a highly linked, central element in the cluster. Ribonucleotide reductase activity is rate limiting for DNA synthesis and contributes to Gemcitabine resistance in human cancers including PDAC (Duxbury et al., 2003; Nakano et al., 2007; Zhan et al., 2021).

### Inhibition of acid-adaptation-regulated genes resensitizes acid-adapted organoids to Gemcitabine

Motivated by these distinct, yet partially overlapping responses, we visualized expression changes of key genes in Gemcitabine and Erlotinib metabolic and -response pathways derived from literature sources (see Methods) using the same log_2_FC values as above (Fig. 4B) from acid adaptation and drug treatments vs respective controls (Fig. 5A).

**Figure 5:**
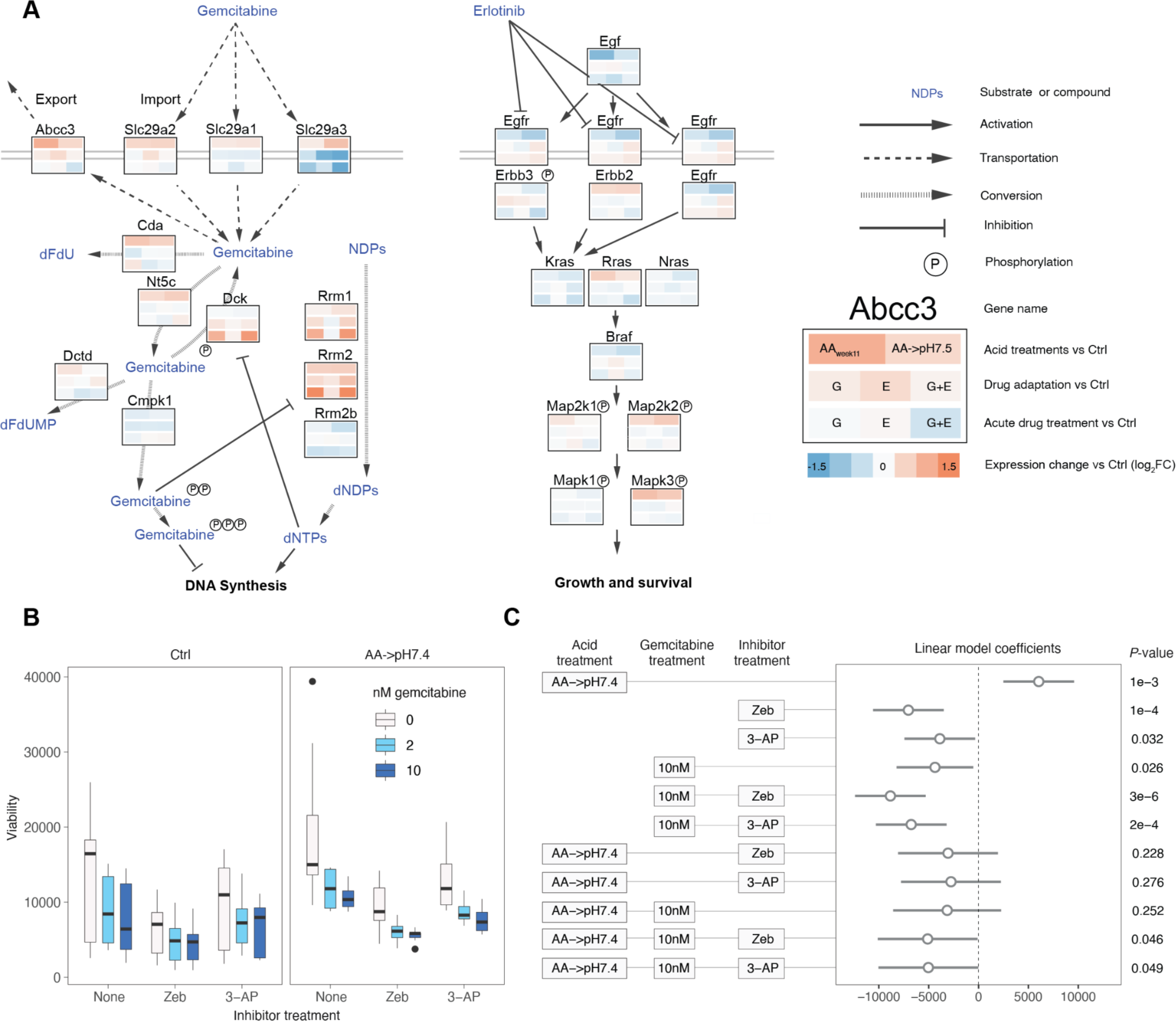
Acid adaptation-induced drug resistance induced through Cda and Rrm2 in mP4 organoids. **A: Expression of key genes in Erlotinib and Gemcitabine pathways following acid adaptation and drug treatments.** Image shows schematic pathways of key Erlotinib and Gemcitabine resistance/metabolic genes and substrates. Lines indicate relationships between genes/proteins as specified in legend. Boxes below gene names show log_2_FC RNA-seq gene expression values (drug or acid adaptation vs. respective control as in Fig. 4; log_2_FC values are capped at +-1.5: see legend). **B: Viability of mP4 organoids following inhibition of Cda and Rrm2.** Boxplots show the viability of Ctrl. and AA→pH7.4 organoids treated with combinations of Gemcitabine and inhibitors as indicated by X axis and color. Boxplots are grouped by acid treatment (indicated on top) and then by inhibitor treatment: Zeb (100 μM Zebularine, which inhibits CDA), 3-AP (5 μM 3-AP, which inhibits RRM2), and no inhibitor control. Color fill represents the dosage of Gemcitabine (0 or 10 nM). **C: Linear regression analysis of the effects of acid adaptation, Gemcitabine, Cda and Rrm2 inhibitors on organoid viability.** The plot to the right visualizes the coefficient estimation from a linear regression of viability as a function of acid adaptation, Gemcitabine concentration, inhibitor treatment and combinations thereof (see Methods). Each row of the plot shows the estimated viability change when pH, Gemcitabine dosage and inhibitor treatment changes as shown by the schematic on the left. The first three rows show the baseline effect of acidosis conditions, gemcitabine, and inhibitors separately; rows 4-6 represent the baseline effect of gemcitabine-inhibitors combinations in ctrl organoids. The last 5 rows show whether gemcitabine-inhibitor combinations cause an additional viability decrease in acid adapted organoids. Estimated effects are represented by hollow circles, with lines indicating the 95% confidence intervals. The *P* value of each estimation is shown to the right. The figure only shows selected combinations of acid adaptation, Gemcitabine concentration, and inhibitor treatment levels: an expanded figure with all levels is shown in Fig. S5A.

Echoing the correlation analysis above, gene expression changes driven by acid adaptation and drug treatment generally differed across the genes in the pathways, indicating that drug resistance is achieved through different trajectories, with a few exceptions. The most striking similarity was the shared upregulation of *Rrm2,* a highly expressed in chemotherapy-resistant cancers (Zhan et al., 2021) and to some degree also the other ribonucleotide reductase subunit, *Rrm1*. This is consistent with our previous analysis where *Rrm2* was also identified as a shared upregulated gene (Fig. 4D). However, other parts of the Gemcitabine pathway showed drastically different expression profiles: nucleoside transporters were upregulated in AA_w11_ and AA→pH7.4 organoids but their expression was unchanged or decreased after drug treatments. Interestingly, Cytidine deaminase (*Cda*), which catalyzes inactivation of Gemcitabine to 2’,2’-difluoro-2’-deoxyuridine (dFdU) (Gusella et al., 2011) and is critical for Gemcitabine resistance in pancreatic cancer(Bjånes et al., 2020), was only upregulated in AA_w11_ and AA→pH7.4 organoids.

Erlotinib is an EGFR tyrosine kinase inhibitor, and resistance to Erlotinib treatment is driven by numerous different genetic changes, including mutations in the receptor itself, constitutively activating KRAS mutations and other activating changes in downstream Ras-Raf-Mek-Erk and PI3K-AKT-mTOR signaling, downregulation of pro-apoptotic Bcl-2 proteins and other death pathways, and upregulation of signaling via alternative receptors (Vaquero et al., 2022). Although there were some divergent changes in EGFR signaling pathways in response to both acid adaptation and drug treatments, the amplitudes of changes were modest, including small changes in the Ras-Raf-Mek-Erk and PI3K-AKT-mTOR signaling pathways.

Because *Cda* expression was increased only in the acid-adapted organoids, and *Rrm2* expression was increased in both acid adapted and drug-resistant organoids, we hypothesized that drug resistance in acid-adapted organoids is partially caused by the high expression of their genes products, and that inhibiting these proteins might resensitize the organoids to chemotherapy treatment. Given the more pronounced changes in the Gemcitabine resistance pathways in the acid-adapted mP4 organoids, we focused our attention on this pathway. mP4 AA→pH7.4 and Ctrl_w11_ organoids were treated with Gemcitabine at low concentrations (2 and 10 nM), in absence or presence of the CDA inhibitor Zebularine (Patki et al., 2021) or the RRM2 inhibitor 3-Aminopyridine-2-carboxaldehyde thiosemicarbazone (3-AP, Triapine) (Finch et al., 1999), and viability was measured after respective treatment. This analysis showed that AA→pH7.4 organoids that were treated by 10 nM Gemcitabine displayed substantially decreased viability if treated by either inhibitor, a pattern that was less strong or not present in Ctrl_w11_ (Fig. 5B). A linear model applied to this data using drug- and inhibitor-untreated Ctrl_w11_ as baseline showed that this was statistically significant (Fig. 5C and Fig. S5A; the interaction between AA→pH7.4, 10 nM Gemcitabine and either inhibitor significantly decreased viability; *P*<0.05 for both inhibitors). Thus, we conclude that the Gemcitabine resistance acquired by mP4 AA→pH7.4 compared to Ctrl_w11_ organoids is at least partially due to the increased *Cda* and *Rrm2* expression levels following acid adaptation.

## DISCUSSION

Poorly perfused tumor regions can give rise to highly aggressive niches characterized by acidosis, hypoxia, or a combination of both (Boedtkjer and Pedersen, 2020) demonstrated in PDAC tumors *in vivo* (Kimbrough et al., 2015; Wojtkowiak et al., 2011). In this study, our main question was whether this would lead to the gain of cancer hallmarks such as increased growth and drug resistance, and whether this would depend on the specific driver mutations. Since it is not possible to control cellular diversity, pH and other environmental factors in tumors *in vivo*, and thereby establish causality, we adapted corresponding *in vitro* organoids models over time from a physiologically normal pH (7.4) to an acidic pH relevant to that of acidic tumor niches. Because they are stem cell based, organoids retain the properties of the native tissue better than cultured cell lines (Czaplinska et al., 2019). We used four organoid lines for adaptation experiments - mN10, corresponding to normal pancreatic cells, and a set of KRAS-mutated (mP) organoids with either intact p53, p53KO or the pH sensitive p53 mutant R273H. To our knowledge, this is the first study to investigate acid adaptation of pancreatic/PDAC organoids and the interaction with driver mutations.

An overall key finding was that acid adaptation and the subsequent return of acid-adapted cells to physiological pH leads to increased organoid viability and drug resistance, but that the gain of these properties is strongly dependent on cells having a wild-type p53 genotype.

The increased drug resistance of the drug treatment-naïve acid-adapted organoids is particularly interesting. Importantly, the observed effects are not due to the well-described biophysical effect of extracellular acidosis on protonation and hence import of weak base drugs (Trédan et al., 2007; Wojtkowiak et al., 2011) because the effects are especially potent in organoids that are returned to physiological pH. Contrary to our initial hypothesis, acid adaptation did not drive the cells to a transcriptional state highly similar to that of cells adapted through continued exposure to Gemcitabine or Erlotinib. Instead, our data shows that the increased drug resistance of acid-adapted organoids likely reflects two components: first, the increase in baseline viability conferred by acid adaptation, and second, changes of expression of specific genes within in cancer drug metabolism pathways, of which most are interestingly not regulated by *de facto* drug adaptation. In particular, we observed that in mP4 cells, two genes associated with Gemcitabine resistance, i.e. the M2 subunit of ribonucleotide reductase, *Rrm2,* and cytidine deaminase, *Cda*, which catalyzes the inactivation of Gemcitabine to dFdU, were upregulated by acid adaption. Inhibition of these genes increased the Gemcitabine sensitivity of acid-adapted mP4 organoids more than that of non-adapted mP4 organoids. Notably, *Cda* expression was not altered when the organoids were adapted to the drugs. Thus, adaptation to these environmentally hostile environments (low pH and drug treatments, respectively) lead to similar phenotypes (decreased sensitivity to Gemcitabine) but the specific gene expression programs through which these phenotypes were reached were largely dissimilar, and while a handful of drug-associated pathways were affected by both adaptations, distinct genes within these pathway responded to respective treatment.

Finally, p53-dependent apoptosis is a well-described mechanism of chemotherapy-induced cancer cell death (Hernández Borrero and El-Deiry, 2021). It is therefore interesting to note that - consistent with our recent findings in Panc02 mouse pancreatic cancer cells (Czaplinska et al., 2022) - the p53KO and -mutant organoids showed a higher baseline viability compared to WT organoids, but lacked the acid-adaptation-induced increase in resistance. In light of previous reports that environmental acidosis elicits p53-dependent cell death, providing a selective growth advantage for cells lacking WT p53 (Williams et al., 1999) the pathway affected may involve downregulation of p53-dependent death mechanisms.

Previous work exploring the impact of the tumor microenvironment on drug resistance has mostly focused on the reduced uptake of weakly basic drugs into cancer cells in acidic environments - the phenomenon of ion trapping (Trédan et al., 2007; Wojtkowiak et al., 2011). However, this clearly does not account for the chemotherapy resistance observed in the present work, which was more pronounced at pH 7.4 than at acidic pH. A series of studies have demonstrated how upregulation of net acid extruding transporters - which also occurs in acid-adapted cells, e.g. (Rolver et al, in press) is associated with decreased sensitivity to chemotherapeutic drugs (Stock and Pedersen, 2017). Very few studies have directly assessed the impact of acid adaptation on chemotherapy resistance. Early work in ovarian cancer and melanoma cell lines showed that adaptation to growth at pH 6.7 was associated with resistance to cisplatin in a manner suggested to involve upregulation of heat shock protein HSP27 (Wachsberger et al., 1997).

In conclusion, our study strongly shows that pancreatic and PDAC organoids can acquire cancer hallmarks through gradual acid adaptation, a change that persists and in many cases increases once the organoids are returned to a physiological pH, contingent on p53 mutations. At present, it is unclear if the acid adaptation phenotypes we observe is primarily an effect of genetic selection, since many cells die during adaptation, or mainly an epigenetic/transcriptional state effect. Relatedly, an important question is whether adapted cells that are returned to normal pH retain the cancer hallmarks over a longer time than studied here. A clinically important question is whether a similar adaptation with similar outcomes occurs in acidic niches in PDAC tumors. If cancer hallmarks can be acquired by cells in acid niches within PDAC tumors and retained even when the extracellular acidity in the tumor microenvironment decreases, e.g. during invasion, improved vascularization, or upon tumor shrinkage during cancer treatment, this may be a mechanism underlying PDAC recurrence.

## METHODS

### Maintenance of 3D organoid cultures

Murine pancreatic cancer organoids (mP4) and normal (“healthy”) pancreatic organoids (mN10) developed from murine pancreatic ducts by the Tuveson group (Boj et al., 2015) were cultured in 24 well plates (BioLite Multidish, Thermo Fisher Scientific, #930186), suspended in 25 or 50 µL domes of Matrigel (Matrigel Growth Factor Reduced, Phenol Red Free, Corning, # 356231). They were fed with 500 µl Mouse Complete Feeding Medium (Boj et al., 2015) that was replaced with fresh medium every 3-4 days. Organoids were split approximately twice a week as needed, as in (Boj et al., 2015) .

### Genetic modifications of mP4 organoids

The mP4 p53KO organoids were established from mP4 organoids using CRISPR/Cas9 technology. Briefly, guide RNA (gRNA) sequence 5’-CACCGTGCACATAACAGACT -3’ targeting exon 4 of the *Trp53* gene was designed using CRISPR Finder (https://wge.stemcell.sanger.ac.uk/find_crisprs). The gRNA was then cloned into a pSpCas9(BB)-2A-puro V2.0 (PX459) plasmid (Addgene #62988). The vector was used for organoid transfection using Lipofectamine® 3000 (Thermo Fisher Scientific) according to the manufacturer’s protocol and (Schwank et al., 2013). The successfully transfected population was selected with puromycin, followed by single cell clone isolation and expansion. *Trp53* KO was verified by western blotting on two independent sets of lysates, performed essentially as in (Andersen et al., 2018). The mP4 p53R273H organoids were developed from mP4 p53KO organoids by transfection with the pTwist-EF1-alpha-puro-Trp53R270H (Twist Biosciences) vector coding murine mutant p53-R270H (corresponding to R273H in human p53). A stable mutant line was established by puromycin selection. Results were verified by western blotting using antibodies against p53 and p21 on two independent sets of lysates.

### Adaptation to low extracellular pH

Organoids were gradually acid adapted from pH 7.4 to pH 6.7 by decreasing pH of the culture medium by approximately Δ= -0.125 pH units every week for nine weeks. Culture medium was acidified by titration with 1 M hydrochloric acid (HCl), and filtered through sterile 0.22 µm filters (Millex®-GP, Millipore express PES membrane Filter Unit, #SLGP033RB). The resulting pH was validated using pH determination strips (Sigma Aldrich) inside the CO_2_ incubator. During acid adaptation, a substantial number of cells in all genotypes died, in particular in the week 5 to week 8 period, presumably reflecting the imposed selection pressure. After reaching the desired pH of 6.7, the organoids were cultured for 2 weeks (week 8-9) at pH 6.7. Afterwards, acid-adapted organoids were split into two fractions: i) cultured at pH 6.7 until week 11 (AA_w11_), and ii) cultured at pH of 7.4 until week 11 (AA→pH7.4) to mimic cancer spread to the bloodstream (pH 7.4) and distant organs e.g. liver (pH 7.0 (Park et al., 1979)) and lungs (pH 7.4 (Effros and Chinard, 1969)).

### CellTiterGlo viability assay

The organoids were dissociated to single cells with Trypsin and counted with Invitrogen Countess II FL instrument. The number of dead cells in a population was evaluated with Trypan Blue (Sigma Aldrich, T8154). 10.000 cells were seeded per 50 ul Matrigel dome as described above. Viability was measured with CellTiterGlo® 3D Reagent (Promega, #G9683).

### Measurement of organoid surface area

For a given condition and PDAC organoid type, 2-3 representative images of organoids were taken per dome. Focal plane was established based on the highest number of organoids present in the dome in focus. Scale bar is 400 µm. The number and sizes of organoids in each image were assessed by an automated organoid detection algorithm Orgaquant (Kassis et al., 2019). The algorithm detects organoids on the focal plane of brightfield images. To optimize detection, we used the pre-trained convolutional neural network model orgaquant_intestinal_v3 but adjusted parameters for the imaging preprocessing (image size=2048 *2048, Contrast=2.5) and prediction (threshold=0.9). The surface area of each organoid was then estimated through the sphere surface area formula, with diameters determined by the average edge lengths of the circumscribed detection box. To avoid counting “half” organoids, any organoid was removed if its center was located within a certain distance to each border (left, right, top, bottom). The distance was defined by 5% of the total width of the picture (5% * 2048).

### Statistical analysis of viability and total organoid surface area as a function of acid adaptation

We used a one-way Anova analysis with the model formula *M ∼ ac + b,* where *M =log(viability)* or M =*log(total organoid surface area per image)*, *ac =* acid adaptation ∈{Ctrl, AA_week11_,AA→pH7.4} and *b* = experimental batch (because both viability measurements and image analysis was made in several experimental batches). One-way Anova tests were made separately for each genotype. In addition, TukeyHSD tests were conducted to compare *M* in Ctrl, AA and AA→pH7.4 within each genotype.

### Statistical analysis of surface organoid surface area distribution as a function of acid adaptation

We estimated log(organoid surface area) for each organoid based on Orgaquant analysis as described above. Estimates were batch corrected using the standardize method in the R package batchtma(Stopsack et al., 2021). Batch corrected data were grouped by genotype and acid adaptation conditions and pooled to obtain conditional distributions. The change of distribution as a function of acid adaptation was then tested by pairwise K-S tests within each genotype.

### RNA isolation and sequencing

Confluent organoid cultures (3-4 50 µl domes/condition) were harvested with cold Cell Recovery Solution (Corning, 354253) and incubated on ice for 30 min to dissolve Matrigel and release the cells. RNA from the organoid pellets was isolated and purified using RNeasy Mini Kit (QIAGEN, 74106) according to the manufacturer’s protocol, including DNase digestion step. RNA concentration was measured using NanoDrop 2000 spectrophotometer and samples were stored at –80°C. RNA was subject to polyA-selected 150 bp paired RNA-seq at BGI, Hongkong, as a paid service.

### RNA-Seq mapping, processing and library quality control

Fastqc (version 0.11.5, https://qubeshub.org/resources/fastqc) and Multiqc (version 1.7(Ewels et al., 2016)) were used to investigate potential RNA-Seq library (paired-end) artifacts. We found a high concentration of 13 base pairs 5’ adapter sequences in all the libraries, which were removed using Trim Galore (0.4.4, https://github.com/FelixKrueger/TrimGalore) with the options ‘–paired –clip_R1 13 –clip_R2 13’. Salmon (version 1.3.0 (Patro et al., 2017) with the parameters ‘-l IU’ was then used to map the adapter-trimmed reads on the mouse transcriptome (M25 GRCm38.p6 Gencode release (Frankish et al., 2021)). The Salmon transcriptome index was built as in https://combine-lab.github.io/alevin-tutorial/2019/selective-alignment/. Libraries contained 46*10^6^ million reads on average, of which on average 89% mapped (SD = 2%, Table S1). Treating the different genotypes RNA-seq libraries made for drug adaptation experiments (see below) separately, we made PCA plots (Fig. S2A) from each log transformed expression matrix. Before computing the principal components, we filtered out lowly expressed genes (using the filterByExpr function from edgeR (Robinson et al., 2010) with settings min.count = 55 and min.total.counts = 40) and regressed out the batch effects with removeBatchEffect (Liu et al., 2015) (4 RNA-Seq batches).

### Differential gene expression analysis

Salmon quantification files were loaded with the tximeta package (Love et al., 2020), to extract the read counts mapped on each transcript. Those were summarized to genes counts with the function summarizeToGene. As we used Limma-voom (Liu et al., 2015) for the differential gene expression analysis, we built a design matrix that would account for the time point, the condition (Ctrl_w1_, AA, and AA→pH7.4), the genotype and the Batch (3 experimental batches). A contrast matrix was then compiled: for each genotype, we defined 4 contrasts, each comparing a treatment condition at a given time point to the Ctrl_w1_ samples. Therefore, each genotype has the following contrasts: AA_w5_ vs. Ctrl_w1_, AA_w8_ vs. Ctrl_w1_, AA_w11_ vs. Ctrl_w1,_ AA→pH7.4 vs. Ctrl_w1._ In the main text and the figures, these contrasts are respectively labelled as “AA_w5_”, “AA_w8_”, “AA_w11_” and “AA→pH7.4”. The filterByExpr function was used to filter out the lowly expressed genes, choosing the parameters “min.count = 55, min.total.counts = 40” so that the mean variance trend plot would satisfy the voom criteria (Fig. S2B). When building the model, we chose to use the function voomWIthQualityWeights to weigh down the influence of low-quality samples (Fig. S2C). The differential gene expression statistics were obtained after successively applying the functions efit, contrast.fit (with the contrast matrix as parameter) and eBayes. Only genes with *FDR* < 0.05 (*P* values were adjusted with Benjamini-Hochberg correction) in at least one contrast were conserved for further analysis. For Fig. 2B, an extra cutoff was added so that each gene must satisfy |log_2_FC| > 0.5 for at least 2 contrasts.

### Gene set enrichment analysis

With the clusterProfiler package(Wu et al., 2021), we conducted a GSEA on each contrast (Fig. 2D), ranking the genes according to their log_2_FC. For a given contrast, any gene that was not filtered out in the differentially expressed gene analysis was used in the GSEA, independently of its *FDR*. As gene sets, we used GO terms, the KEGG pathways (Kanehisa et al., 2021), and the Msig database(Liberzon et al., 2015) (category 2) . For Fig. 2D, genes sets of interest were selected and normalized enrichment scores were capped at –2 and 2. The displayed Msig terms’ names were changed to more succinct names for clarity (Table S2).

### Organoid drug treatments

To illustrate the effects of acid adaptation on organoid drug resistance, we treated Ctrl_w11_, AA and AA→7.4 organoids with different concentrations (10 nM, 25 nM) of Erlotinib (Sigma, #SML2156), Gemcitabine (Sigma, #G6423) or a combination of both. Organoid viability and surface area were measured as described above. Organoids were cultured for 5 days in organoid feeding medium containing 10 or 25 nM of Gemcitabine (Sigma, #G6423) or Erlotinib (Sigma, #SML2156), or a combination of both at appropriate pH. After 5 days, brightfield images were acquired on an Olympus IX83 microscope, using a SP 10 objective and CellSens Dimension software. Organoid viability was developed using CellTiterGlo reagent (Cat #G9683, Promega) and detected using FluoStar Optima Plate Reader (21999, BMG LabTech).

### Statistical analysis of viability and total organoid surface area as a function of drug treatment

We used a one-way Anova analysis with the model formula *M ∼ dt + b, (*where *M =log(viability),* as defined above or M= *log(mean(total organoid surface area per image),* as defined above), *dt =* drug treatment ∈{Untreated, E10, G10, E+G10} and *b* = experimental batch (because both viability measurements and image analysis was made in several experimental batches). One-way tests were made separately for each combination of acid adaptation condition (Ctrl, AA_w11_ and AA→pH7.4) and genotype. To assess the effect of acid adaptation condition and drug treatment and their possible interaction on viability and total surface area, for each genotype and drug type (E, G, E+G), we use a linear model with the formula *M ∼ ac + dt + ac*dt + b*, where *M* was defined as above, *ac* = acid adaptation condition ∈{Ctrl, AA, AA→pH7.4} with Ctrl. as the baseline, *dt* = drug treatment ∈{untreated, drug 10 nM, drug 25 nM} where drug ∈ {E,G, E+G} and untreated is used as baseline and *b* is the experimental batch as above. In Fig. S3 C-D, for the drug dosage X ∈{10, 25}, “Drug XnM” models the effect of *dt* while “AA x D.X” and “AA→pH7.4 x D.X” model the interaction of *ac* and *dt* (*ac*dt)* for AA or AA→pH7.4 acid adaptation conditions, respectively. The model was fitted using the lm function which calculates *t*-statistics and associated *P*-values. Distribution of log(organoid surface area) across different genotypes, drug treatments and acid adaptation conditions were analyzed based on the batch corrected data and Kolmogorov-Smirov tests as described above.

### Drug adaptation of organoids and subsequent analysis

PDAC organoids (mP4 genotype) were gradually adapted to growth in medium containing 10 nM of drugs (Erlotinib, Gemcitabine or combination), by increasing drug concentrations by 1 nM/week over a period of 12 weeks. RNA samples were collected for RNA-sequencing from week 1 and week 12, as above. For confirmation of resistance development (Fig. S4A-C), 5,000 cells were seeded in Matrigel domes in a 48-well-plate (LABSOLUTE®, Art.N 7696793, Th. Geyer GmbH & Co.KG) with medium containing increasing concentrations (0, 5, 10, 20, 50 and 100 nM) of Gemcitabine, Erlotinib, or a combination of both. Viability was determined after 5 days using CellTiterGlo reagent (Cat #G9683, Promega) and detected using FluoStar Optima Plate Reader (21999, BMG LabTech). RNA-seq analysis on these samples was performed in biological triplicates as described above. Apart from the filtering (performed using filterByExpr with the default parameters), the RNA-seq analysis was performed as for Fig. 2. We defined 8 contrast : week 12 E, G and E+G drug adapted or acutely treated samples were compared to Ctrl_w12_ from the drug adaptation experiment while the AA_w11_ and the AApH→7,4 samples were compared to Ctrl_w11_ in the acid adaptation experiment. For each contrast defined above, we calculated corresponding log_2_FC values (treatment vs. control), and then calculated the Pearson correlation between drug treatment and acid treatment log_2_FC values in a pairwise manner.

### Permutation tests

For each pairwise comparison of treatments vs. control log_2_FC values from RNA-seq data described above, genes with log_2_FC > 0.25 in both treatments were defined as jointly upregulated. We define the ratio of jointly upregulated genes vs. all other genes as the observed joint upregulated ratio up_observed_. To assess whether this ratio was higher or lower than expected given the log_2_FC distributions, we shuffled the gene labels within each log_2_FC vector, and then calculated the ratio of jointly upregulated genes in the same way as above. We repeated this 10^5^ times, and calculated the mean of the resulting ratios. We define this as the expected joint upregulated ratio, up_expected_. The observed vs. expected ratio was then expressed as log_2_(up_observed_/up_expected_). The same procedure was used for jointly down regulated genes, defined as log_2_FC < -0.25 in both treatments.

### Visualization of shared responsive genes in acid and drug treatments

The String database version 11.5 (Szklarczyk et al., 2019) was used to visualize the selected shared up- and down-regulated genes in acid adaptation and drug treatments defined as above. The minimum required interaction score was set to high confidence (0.700) and disconnected nodes in the network were not shown. Selected terms were color-coded based on the functional enrichment analysis within the STRING database web interface.

### Pathway visualization

We visualized expression profiles of PDAC- or drug resistance related-genes with Cytoscape software. Genes are selected from well characterized pathways from literature or the Wikipathway database (Martens et al., 2021), in particular the Gemcitabine (Alvarellos et al., 2014) and EGFR signaling pathways (https://www.wikipathways.org/index.php/Pathway:WP4806).

### Zebularine and 3-AP inhibition experiment

For a given PDAC organoid type in three biological replicates, 5.000 cells were seeded per 25 µl Matrigel dome in a 48-well-plate (LABSOLUTE®, Art.N 7696793, Th. Geyer GmbH & Co.KG) with medium containing increasing concentrations of Zebularine (0, 100 μM; 2BScientific, #3690-10-6), 3- AP (0, 5 μM; Sigma, #SML0568), or Gemcitabine (0, 2, 10 nM; Sigma, #G6423) or combination of the three. Viability of organoids was determined after 3 days using CellTiterGlo reagent (Cat #G9683, Promega) and detected using a FluoStar Optima Plate Reader (21999, BMG LabTech).

### Linear regression analysis for inhibitor assays

We fitted a linear model for organoids viability as a function of acidosis condition, Gemcitabine treatment, and inhibitors (Zebularine, 3-AP): V ∼ ac + gt_in + ac * gt_in + b, where V was viability, ac = acid adaptation condition ∈{Ctrl, AA→pH7.4}, gt_in = combination of Gemcitabine treatment and inhibitors ∈{untreated, Gemcitabine 2 nM without Inhibitors, Gemcitabine 2 nM with Zebularine 100μM, Gemcitabine 2 nM with 3-AP 5 μM, Gemcitabine 10 nM without Inhibitors, Gemcitabine 10 nM with Zebularine 100 μM, Gemcitabine 10 nM with 3-AP 5 μM}, b = experiment batch. The R function lm() was used to fit the model, estimate the coefficients and calculate *P* values.

### Statistical software

R (version 4, https://www.R-project.org/.) was used for all analyses. Visualization was done using ggplot(Wickham, 2009), ggpubR (https://CRAN.R-project.org/package=ggpubr), pheatmap (https://CRAN.R-project.org/package=pheatmap) and clusterprofiler (Wu et al., 2021) packages. Batch correction for distribution analysis was done using the batchtma (Stopsack et al., 2021) package.

## DATA AVAILABILITY

RNA-seq data from the study has been uploaded to the GEO database with accession number GSE218178. This is composed of two sub series: acid adaptation experiment GSE218173 (security token for referees: spkxgokqdrkhxgz) and drug adaptation experiment GSE218177 (security token for referees: qhmjuouczjwxdcx).

## ACKNOWLEDGEMENTS

We gratefully acknowledge Signe Meng, Nanditha Prasad, and Christian Dalager Vaagensø, University of Copenhagen, DK, for excellent technical assistance, and Luis A. Pardo, Max Planck Institute for Interdisciplinary Sciences, Göttingen, DE, for fruitful discussions and introduction to organoid culture.

## AUTHOR CONTRIBUTIONS

Conceptualization: ASa, SFP. Supervision ASa, SFP, RA, DC. Experimental design: ASa, SFP, DC. Experimental work: DC; RI, HBA. Data analysis and figure preparation: ASt, JY, YD, DC, RI, HBA, AS. First draft: AS, SFP. Paper writing and editing: SFP, ASa, ASt, JY, YD, DC, RI, HBA. Funding acquisition: SFP, AS, DC.

## SUPPLEMENTARY FIGURES WITH LEGENDS

**Figure S1:**
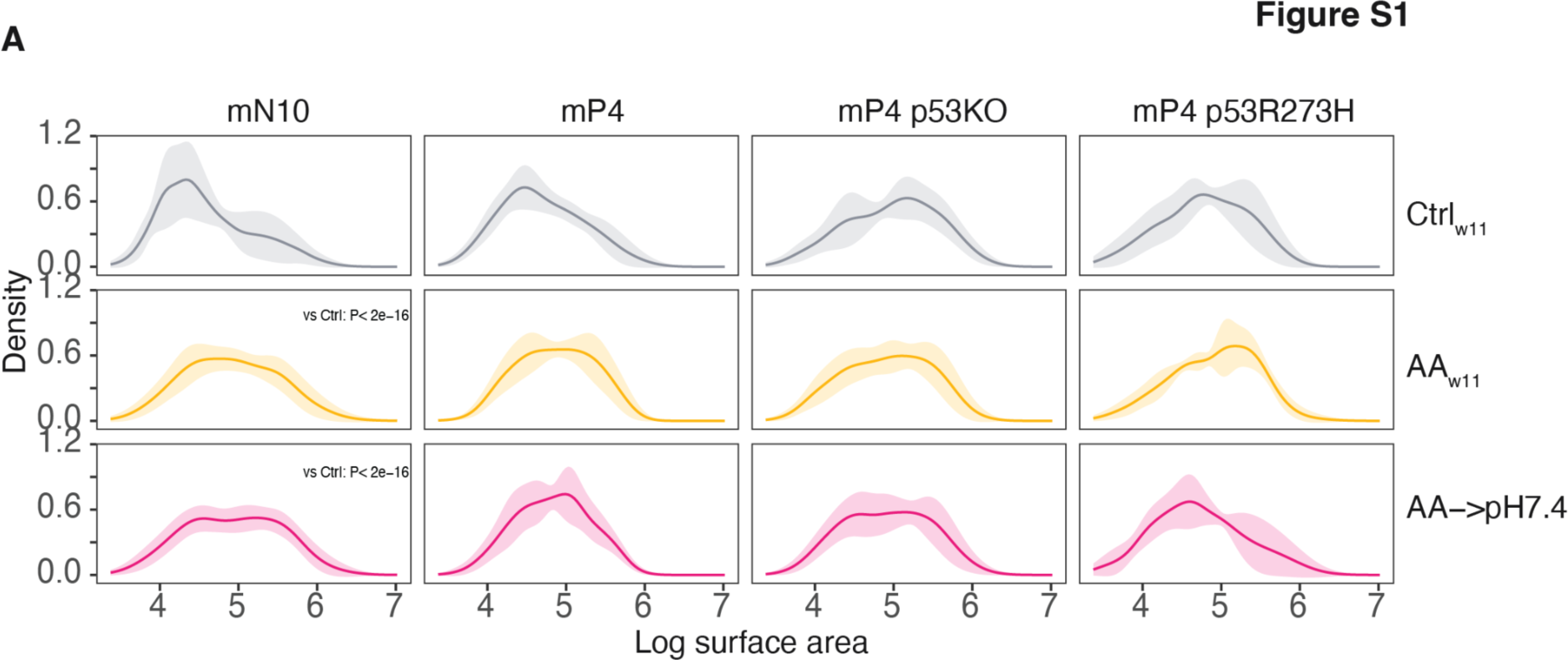
Organoid size distribution estimates. This supplementary figure expands Fig. 1. **A: Size distributions of organoids following acid adaptation** Each subplot shows the distribution of organoid sizes (log scale) for a given acid adaptation state (rows) and genotype (columns). The data is the same as in Fig. 1F, but with color fills that show the mean ± standard deviation ranges of local density, assessed across replicates.

**Figure S2:**
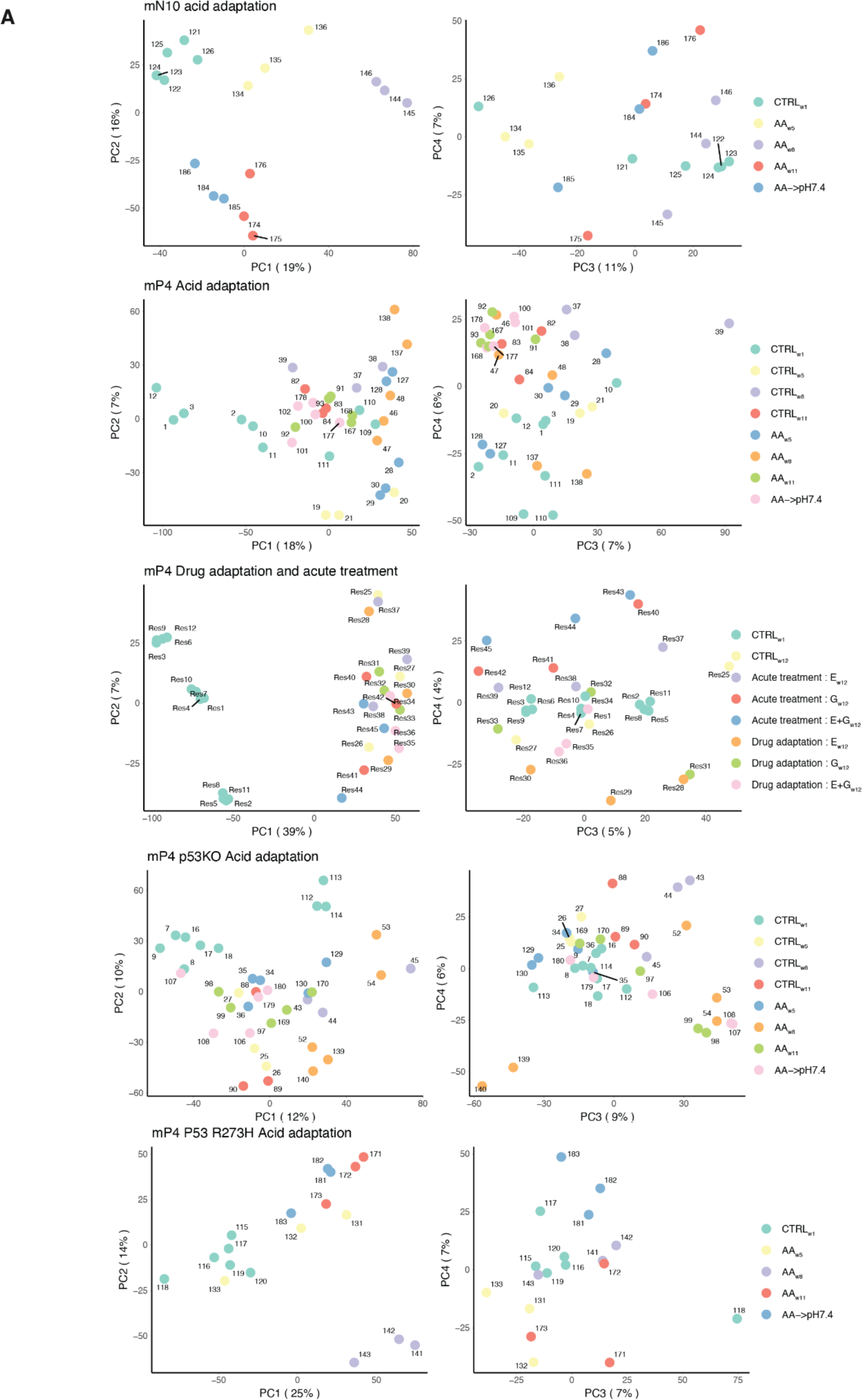
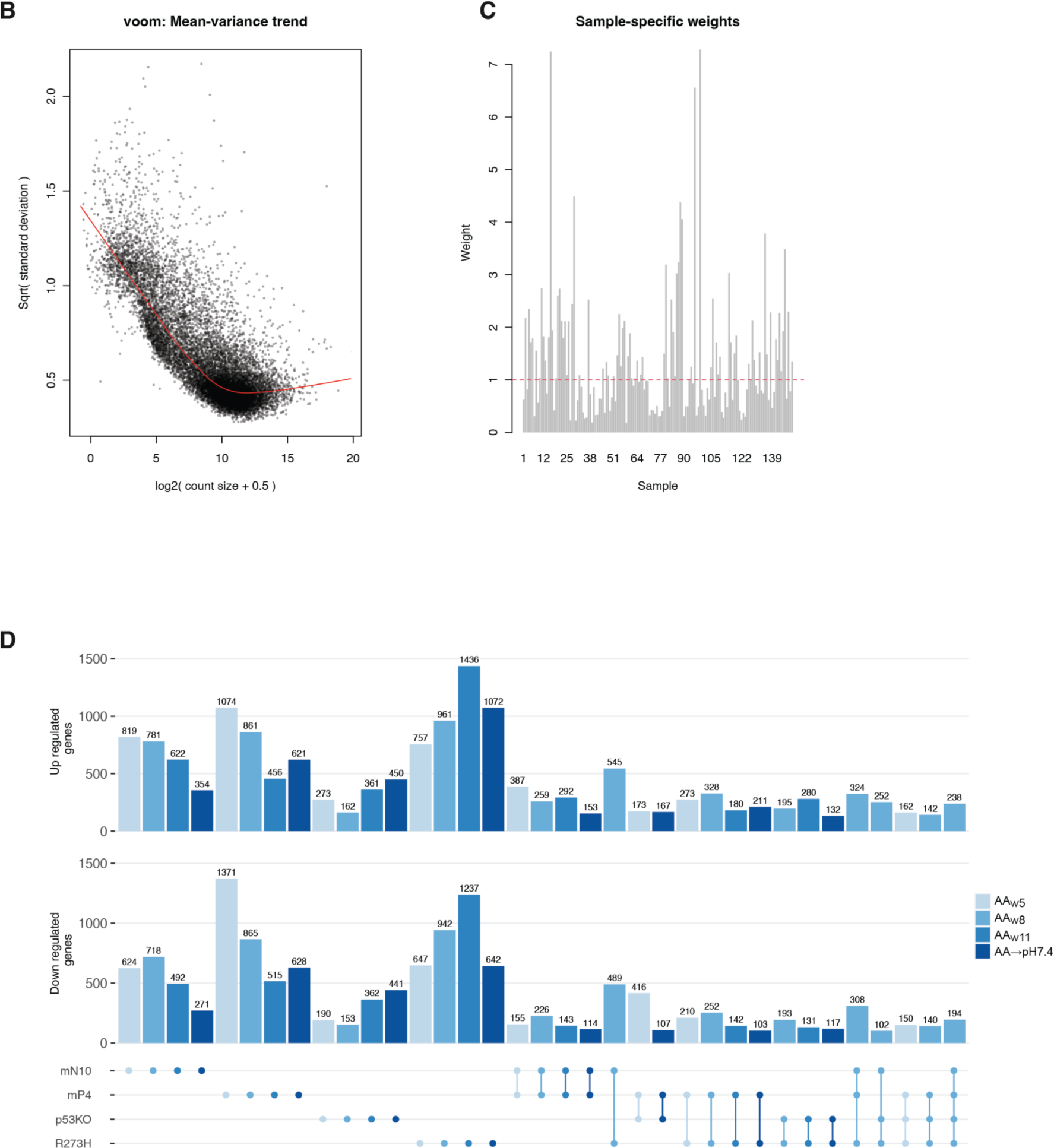
RNA-seq data analysis. This supplementary figure expands Fig. 2. **A: Principal component analysis (PCA) of all RNA-seq data.** Each subplot shows PC1 vs. PC2 and PC3 vs. PC4, analyzing each experiment type and genotype separately. Dots indicate RNA-seq libraries, colored by treatment type. Dot labels correspond to library IDs in Table S1. Data has been batch-corrected before analysis, as described in Methods. Percentages on axes indicate percent of variance explained. For mP4 cells, PCAs for drug-treated organoids are also shown separately (data used in Fig. 4-5). **B-C: Diagnostic plots for voom-based weighing of samples.** The B panel shows relation between the mean(expression) and the variance(expression) for each gene across all libraries after filtering out lowly expressed genes (as described in *Differential gene expression analysis* in Methods). The red curve shows the voom mean-variance trend. The C panel shows the weights attributed by the *voomWithQualityWeights* function to each library. **D: Overlap between differentially expressed genes in the acid adaptation time course.** This upset-plot expands Fig. 2C. Y axis shows the number of up- or down -regulated genes in a given gene set (see dot plot at the bottom) . Bar color indicates acid adaptation stage; numbers on bars is the number of differentially expressed genes that overlap in the given gene set. Sets of differentially expressed genes are shown at the bottom as dots. If genes were differentially expressed in several genotypes, the dots are connected. Overlaps or sets with less than 100 genes are not shown.

**Figure S3.**
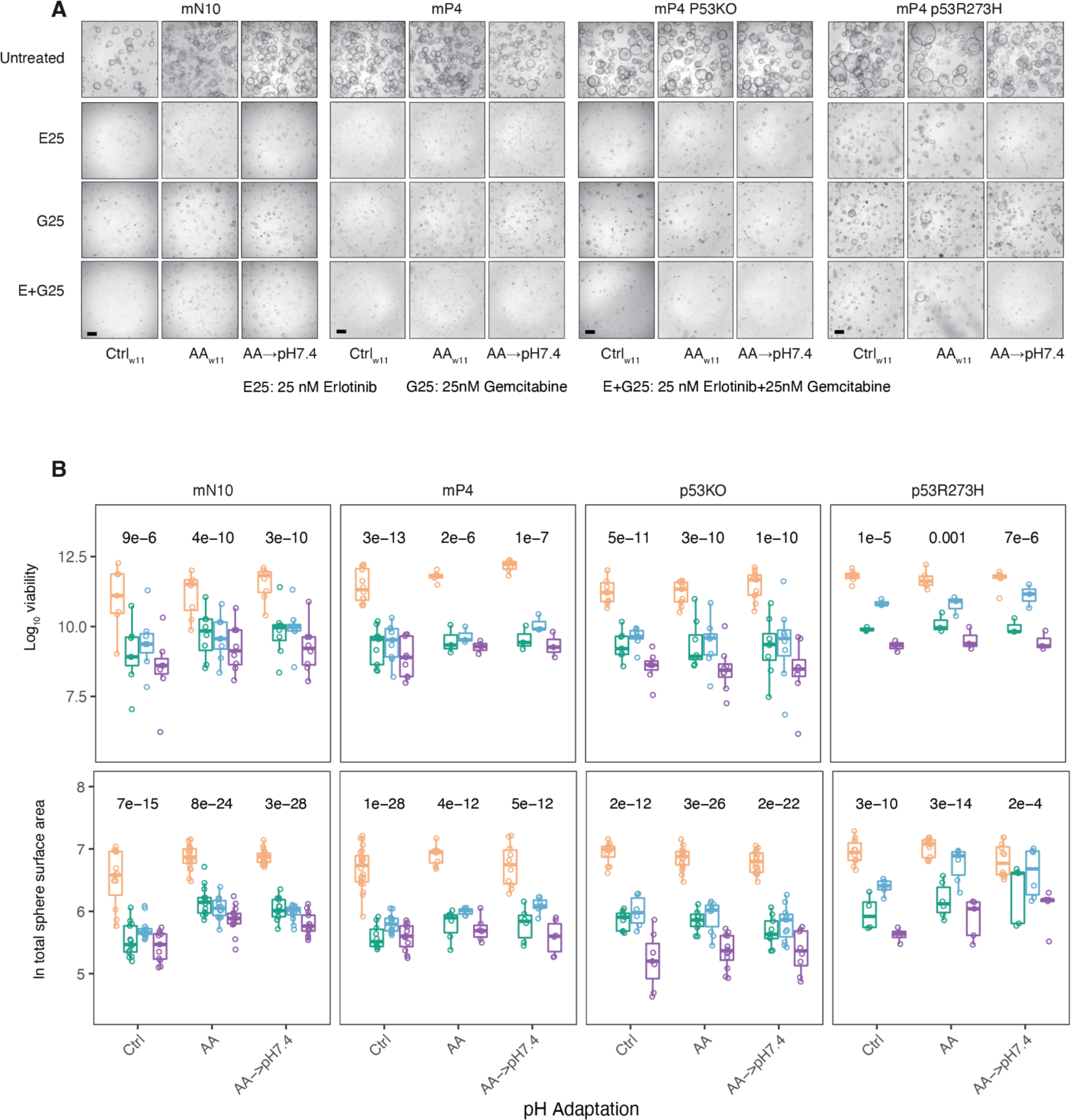
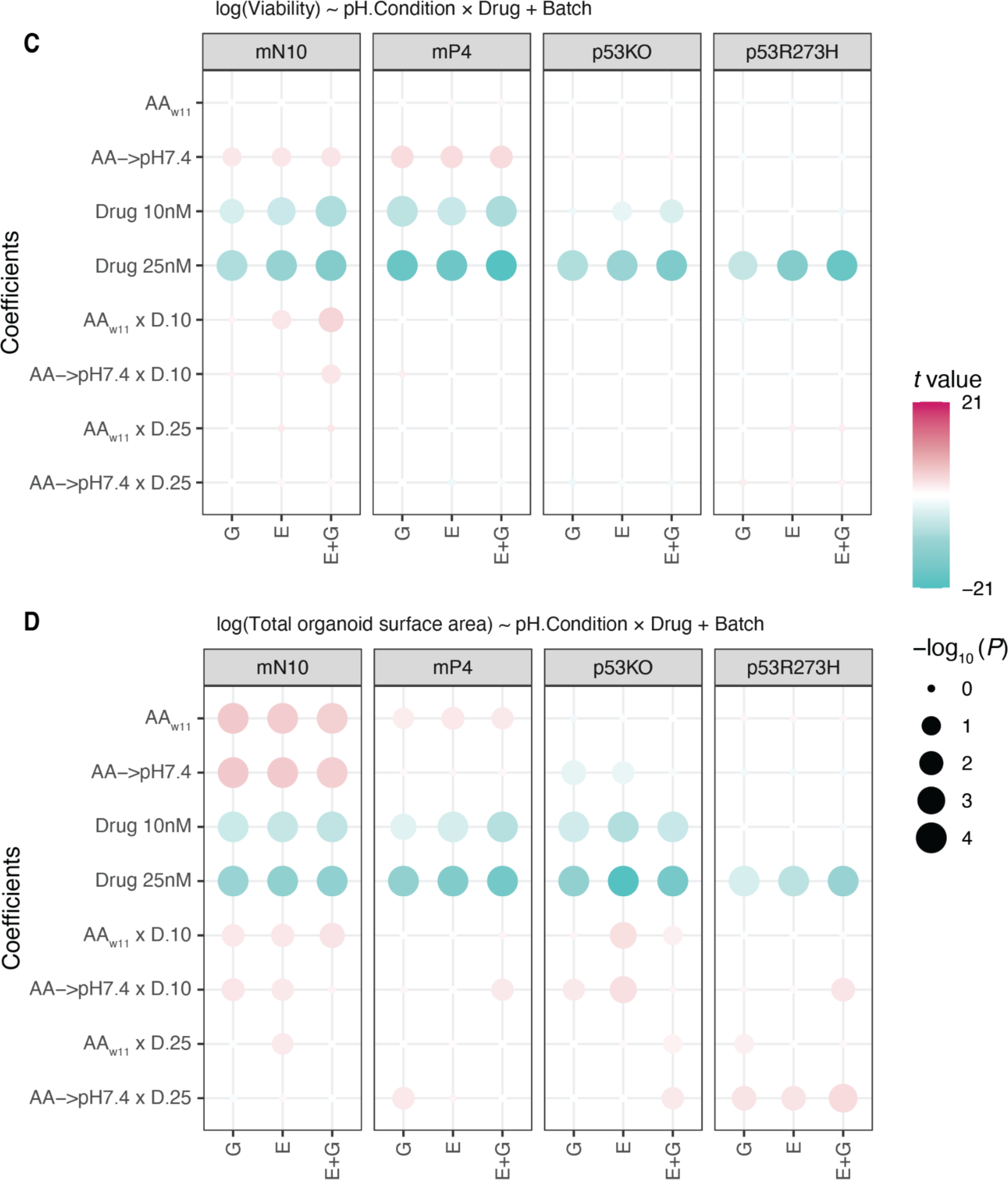
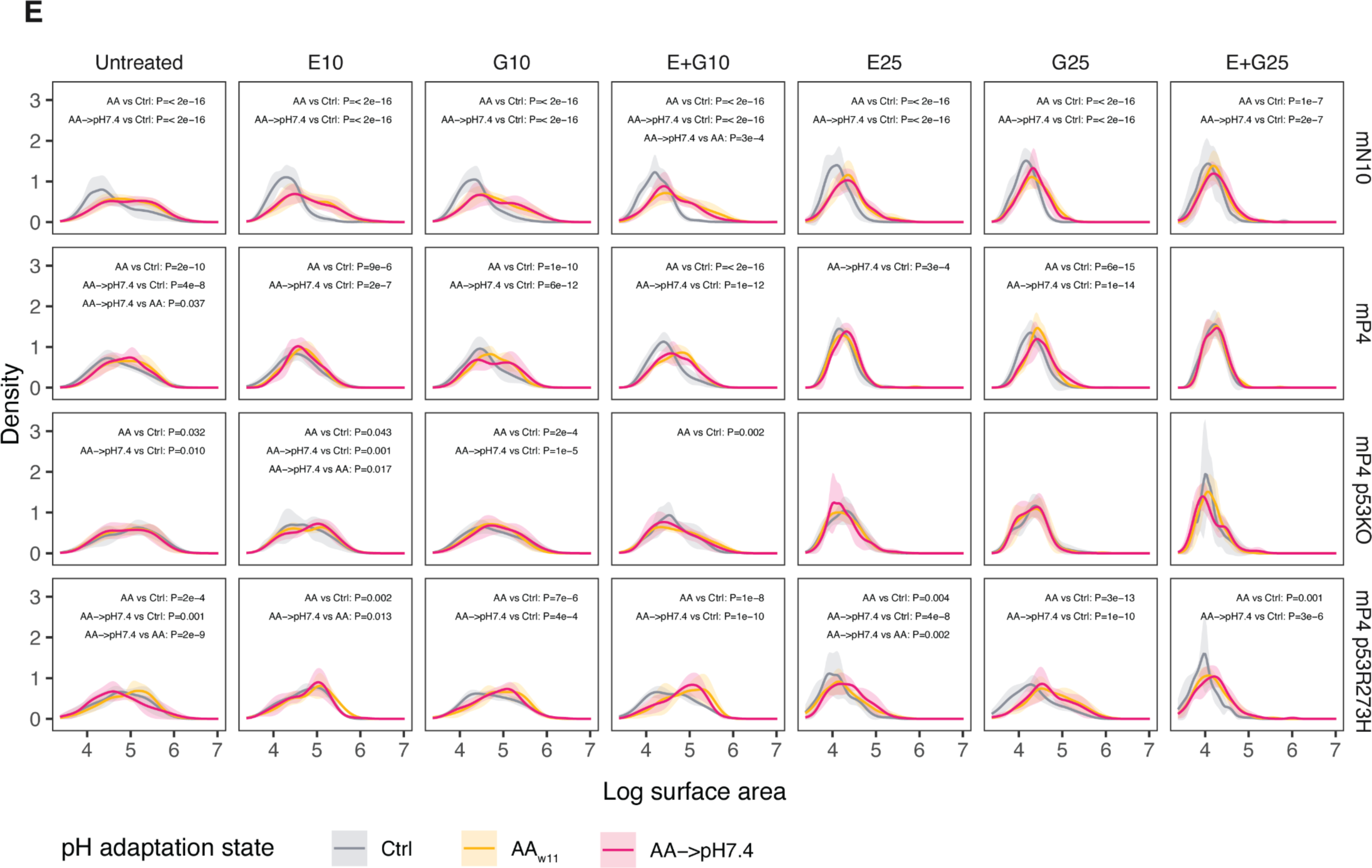
Analysis of relation between acid adaptation and drug treatment response. This supplementary figure expands Fig. 3. **A: Representative images of drug-treated and/or acid-adapted organoids.** The plot is organized as in Fig. 3A, but images show 25 nM drug treatments. Note that the ‘Untreated’ row is identical to that of Fig. 3A. Scale bar: 400 μm. **B: Distributions of viability and total organoid surface area following drug treatments.** As in Fig. 3B, but showing 25 nM drug treatments. Note the drug-untreated organoids boxplots (yellow) are also shown in Fig. 1E and 3A. **C: Regression coefficients from linear models predicting log(viability) by acid adaptation, drug treatment and their interaction.** Based on data shown in Fig. 3B, log (viability) was regressed by pH adaptation, drug treatment and interaction of those two factors, separately for each genotype and drug type (see Methods). Each regression model is shown on one column, where effect terms are shown on rows (AA: Acid adaptation effect on log viability in untreated organoids; AA→pH7.4: AA→pH7.4 effect in untreated organoids; Drug 10 nM: effect of 10 nM of indicated drug in Ctrl_w11_ organoids; AA x D.10: the different effect of 10 nM drug in AA organoids compared to Ctrl_w11_, etc.) Coefficient *t* values and -log_10_ *P*-values are represented by dot colors and dot sizes, respectively. **D: Regression coefficients linear models predicting log (total organoid surface area) by acid adaptation, drug treatment and their interaction.** The plot is organized as in Fig. S3C, but the regression is based on log (total organoid surface area). **E: Distributions of individual organoid surface area.** The plot is organized as in Fig. 1F, but includes measures from all drug treatments. Note that the untreated columns correspond to the data in Fig. 1F. *P* values indicate Kolmogorov-Smirnov tests between distributions, but only significant *P* values (*P*<0.05) are shown.

**Figure S4:**
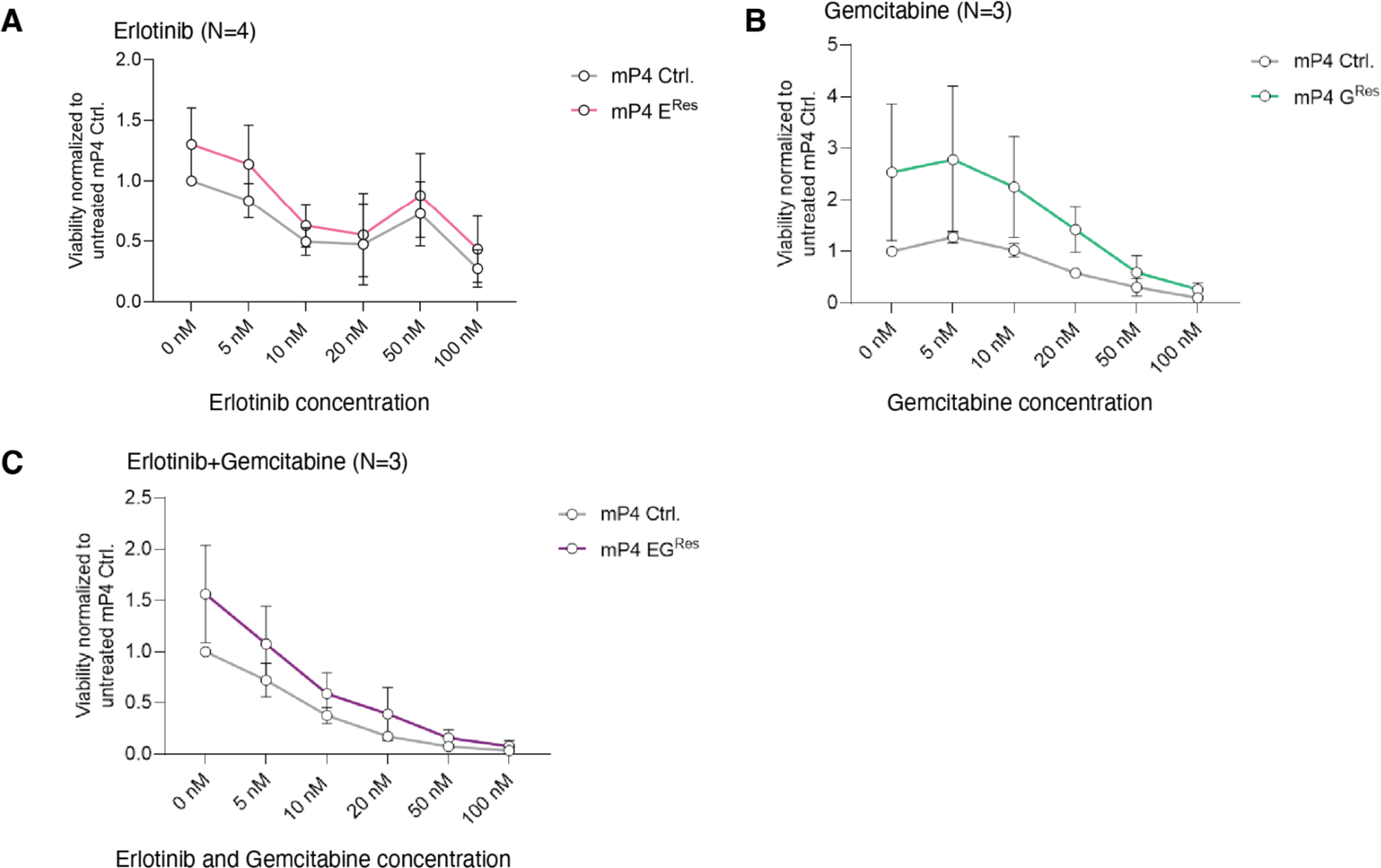
Verification of development of drug resistance. This supplementary figure expands Fig. 4. **A: Erlotnib resistance of Erlotinib-adapted mP4 organoids.** mP4 organoids that were subject to gradual Erlotinib adaptation (mP4E^Res^, also see Fig. 4A) and control mP4 organoids (mP4 Ctrl) were treated with increasing Erlotinib concentrations (X axis), followed by viability assays: Y axis show average viability as defined in Fig. 1E, but normalized to untreated mP4 Ctrl organoids. Error bars show standard error of the mean, number of replicates at concentration and experimental group is indicated on top. **B: Gemcitabine resistance of Gemcitabine-adapted mP4 organoids.** mP4 organoids that were subject to gradual Gemcitabine adaptation (mP4G^Res^, also see Fig. 4A) and control mP4 organoids (mP4 Ctrl) were treated with increasing Gemcitabine concentrations (X axis), followed by viability assays: Y axis as in panel A. **C: Combined Erlotnib+Gemcitabine resistance of Erlotnib+Gemcitabine-adapted mP4 organoids.** mP4 organoids that were subject to gradual Gemcitabine+Erlotinib adaptation (mP4EG^Res^, also see Fig. 4A) and control mP4 organoids (mP4 Ctrl) were treated with increasing Erlotnib+Gemcitabine concentrations (X axis), followed by viability assays: Y axis as in panel A.

**Figure S5:**
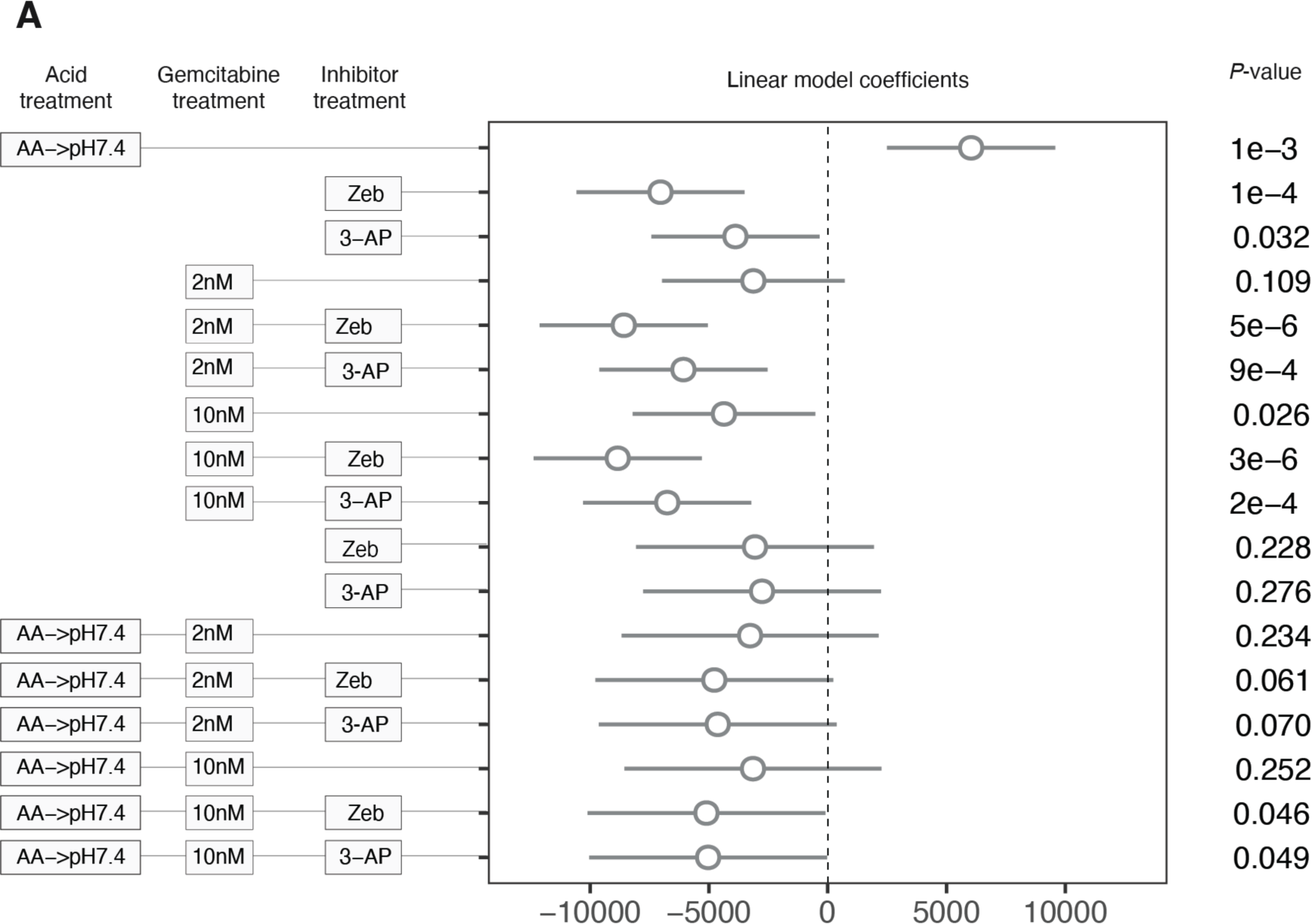
Linear regression analysis of the effects of acid adaptation, Gemcitabine, Cda and Rrm2 inhibitors on organoid viability. This supplementary figure expands Fig. 5. **A: Expanded linear regression analysis.** The figure is organized as in Fig. 5B, but includes coefficients for all Gemcitabine dosages: 0, 2 nM and 10 nM.

## SUPPLEMENTARY TABLES

**Table S1:** List of all RNA-seq samples, their group, batch, library name, genotype, time point (in weeks), pH, and library statistics, including total number of reads and mapping rate. Available at <u>10.6084/m9.figshare.21716540</u>

**Table S2:** List of abbreviated and full gene set names used in Fig. 2D. Available at <u>10.6084/m9.figshare.21716567</u>

